# α-Actinin-3 deficiency protects from the effects of acute cold exposure through altered skeletal muscle Ca^2+^ and OXPHOS signaling

**DOI:** 10.1101/2024.08.14.605938

**Authors:** Peter J Houweling, Chrystal F Tiong, Leonit Kiriaev, Roberto Díaz-Peña, Patrick K. Albers, Harrison Wood, Victoria Wyckelsma, Tegan Stait, Tomas Venckunas, Pedro L. Valenzuela, Adrián Castillo-García, Marius Brazaitis, Yemima Berman, Jane T. Seto, Håkan Westerblad, Monkol Lek, David Thorburn, Alejandro Lucia, Stewart I Head, Kathryn N North

## Abstract

A nonsense polymorphism in the *ACTN3* gene (R577X, rs1815739) results in the loss of the fast skeletal muscle fiber protein α-actinin-3 in an estimated 1.5 billion humans worldwide. Homozygosity for this common polymorphism (*ACTN3* 577XX) does not cause disease but is detrimental to sprint performance in elite athletes. Recently, we reported that *ACTN3* 577*XX* humans and *Actn3* knockout mice show improved cold tolerance. This was not due to an increase in brown adipose tissue activity or a greater muscle shivering response, but was associated with an increased abundance of slow myosin heavy chain muscle fibers in XX individuals, which in turn increased muscle tone to improve cold tolerance.

Using the *Actn3* knockout mouse we have now analyzed the molecular and physiological impacts of acute cold exposure (4°C, for 5 hours) on skeletal muscle. In addition, we used statistical inference of publicly available archaic (Neanderthal and Denisovan) and modern human DNA data to estimate the age of the *ACTN3* X-allele and to identify a novel haplotype associated with this variant that provides support for genetic selection in some populations of modern humans.

Our results indicate that skeletal muscle of *Actn3* knockout mice exhibit higher mitochondrial oxidative phosphorylation abundance and activity than wild-type controls. Following acute cold exposure, we identified reductions in gene ontology pathways associated with metabolism and protein breakdown, which impact muscle mass loss and improved muscle fatigue resistance. Together this provides further evidence for improved cold tolerance in *Actn3* knockout mice and *ACTN3* 577XX individuals. Furthermore, we confirm that the X-allele first appeared in modern humans ∼135,000 years ago, with this variant increasing in frequency in European and Asian populations from as early as 42,000 years ago, which aligns with modern human migration out of Africa. Taken together these data provide additional molecular and functional evidence for a protective advantage of α-actinin-3 deficiency following cold exposure, highlighting the importance of skeletal muscle in mammalian thermogenesis and its potential role in the evolution of modern-day humans.

## Introduction

The *ACTN3* gene encodes the skeletal muscle protein α-actinin-3, which is expressed in fast, glycolytic fibers. An estimated 16% of people worldwide (∼1.5 billion individuals) are α-actinin-3–deficient due to homozygosity for a nonsense polymorphism (rs1815739, *ACTN3* 577XX) in the *ACTN3* gene [1].

α-Actinin-3 deficiency does not cause disease per se but can be detrimental to sprint and power performance, in elite athletes [2, 3] and the general population [4–7]. Conversely, there is a positive association between α-actinin-3 deficiency and endurance performance [3, 8], adaptive response to endurance training [9] and maximum aerobic capacity [10–12].

Intriguingly, the frequency of the *ACTN3* 577X null allele differs between human populations; the X-allele is present in <10% of African and >50% of European and Asian populations. Previous work has suggested that the 577X null allele underwent strong, recent positive selection as modern humans migrated out of Africa into the Northern Hemisphere [13], and that the *ACTN3* 577XX genotype has increased in frequency along a global latitudinal gradient, with the null genotype more common in populations inhabiting geographic regions with lower mean annual temperatures [14]. Amorin and colleagues replicated these findings in a separate population, providing additional support for the selective advantage of α-actinin-3 deficiency as modern humans migrated from Africa 40,000–60,000 years ago [15]. More recently, this work was questioned by a study suggesting that the X-allele frequency differences are driven by genetic drift rather than positive selection [16].

Much of the mechanistic understanding of α-actinin-3 deficiency comes from studies using the *Actn3* KO mouse model. *Actn3* KO mice mimic many of the phenotypes seen in α- actinin-3–deficient humans, including lower muscle mass and strength, as well as higher endurance capacity, fatigue resistance and response to exercise training (as reviewed in [17]). The muscles of *Actn3* KO mice does not exhibit a different fiber type distribution compared to wild-type (WT) mice, but their ‘fast’ (i.e., MyHC IIb) fibers, where α-actinin-3 is predominantly expressed, display a shift towards a slower muscle phenotype, including lower diameter [18], higher oxidative metabolism [2, 13, 18, 19], slower muscle relaxation following contraction [20, 21], and faster recovery from fatigue [2, 20]. *Actn3* KO mouse and *ACTN3* 577XX human skeletal muscles also show higher glycogen storage (due to a lower glycogen phosphorylase activity) [19], higher calcineurin activity [9] and altered Ca^2+^ handling [21] that together promote greater oxidative metabolism associated with the ‘slower’ muscle phenotype. We now know that the absence of α-actinin-3 influences an array of structural, signaling, metabolic and protein synthesis/breakdown pathways that together modify muscle function and performance (reviewed in [22] and [17]).

We have previously reported that α-actinin-3–deficient humans and *Actn3* knockout (KO) mice maintain a higher core body temperature during an acute cold challenge. At a physiological level, cold-exposed *ACTN3* 577XX humans activate effective heat generation by increased muscle tone rather than overt shivering [23]. Furthermore, muscles collected prior to cold exposure show an overall lower abundance of fast MyHC muscle isoforms, and a compensatory higher expression of slow type I MyHC protein compared to α-actinin-3 577RR expressing muscles. . Using the *Actn3* KO mice, we showed that the improvement in core body temperature is not due to an increase in brown adipose tissue (BAT) activity. Taken together we concluded that the loss of α-actinin-3 provides a protective benefit during cold exposure due to changes in the functional properties of *ACTN3* 577XX skeletal muscle [23]. However, muscle biopsies were not available post cold exposure for further molecular phenotyping.

In this study, we aimed to further explore the effects of cold exposure by assessing mice housed under acute cold (4°C, for 5 hours) and thermoneutral (30°C) conditions to provide a detailed investigation into the molecular and physiological consequences in acute cold exposed skeletal muscle of α-actinin-3 deficient. We hypothesized that *Actn3* KO muscles would be more resistant to the effects of cold exposure and therefore provide further evidence to support the potential role of α-actinin-3 during the evolution of modern humans as well as the importance of skeletal muscle thermogenesis during cold exposure.

Using the *Actn3* KO mouse we describe *Actn3* genotype-specific changes in key metabolic and mitochondrial oxidative phosphorylation (OXPHOS) pathways as well as altered Ca^2+^ sensitivity, which culminate in improved thermoregulation, a reduction in muscle mass loss and improved muscle function following acute cold exposure. Together these data confirm that α-actinin-3 deficiency enhances skeletal muscle thermogenesis to protect XX/KO individuals from the effects of acute cold stress, which would be beneficial for survival in cold environments. This work provides additional molecular and physiological evidence that highlights skeletal muscle as a critical thermogenic tissue.

## Results

### RNA sequencing supports reduced activation of key thermogenic, Ca^2+^ and metabolic signaling pathways in cold-exposed *Actn3* KO skeletal muscles

Our previously published work demonstrated that cold-exposed α-actinin-3–deficient humans (*ACTN3* XX) and *Actn3* KO mice maintain a higher core body temperature (>35.5°C), compared to those that express α-actinin-3 (*ACTN3* 577RR humans and WT mice [23]).

To further explore the acute effects of altered ambient temperature and define the broader molecular consequences of cold exposure, we performed RNA-sequencing (RNA-seq) on the quadriceps skeletal muscles of WT and *Actn3* KO mice housed at different ambient temperatures. Firstly, a principal component (PC) analysis showed that samples were separated based on both temperature (PC1) and genotype (PC2, Fig. 1A). This is confirmed by a hierarchical clustered heat map that linked samples based on temperature and genotype, with the largest difference between the thermoneutral and cold-exposed muscle samples (Fig. 1B). A total of 4,076 differentially expressed genes were observed (false discovery rate (FDR) p<0.05, Fig. 1C). After 5 hours of cold exposure, 38 genes (28 down-regulated and 10 up- regulated) were differentially expressed in KO mice compared to WT (FDR p<0.05, Fig. 1D). The top 10 genes, included two genes (*AW551984* and TNF receptor associated factor 3 (*Traf3*)) whose expression was upregulated in KO muscles and downregulated in WT, and eight genes (ankyrin repeat domain containing 26 (*Ankrd26*), angiotensinogen (*Agt*), GABA type A receptor associated protein like 1 (*Gabarabpl1*), GTP cyclohydrolase I (*Gch1*), H3 histone family member 3B (*H3f3b*), Ras related dexamethasone induced 1 (*Rasd1*), *Rinagl1*, and zinc finger with KRAB and SCAN domains 14 (*Zkscan14*)) that are downregulated in KO compared to WT (Fig. 1E). Analysis of these genes (FDR P<0.05) using Metascape [24] highlighted gene ontology pathways that are differentially expressed in KO muscles following cold exposure, including *‘muscle cell differentiation’*, ‘*regulation of growth*’, *‘calcium regulation’* and *‘metabolism’*(Fig. 1F). Gene set enrichment analysis (GSEA) of the Kyoto Encyclopedia of Genes and Genomes (KEGG) pathways further identified key metabolic terms that differ with *Actn3* genotype and temperature (Fig. 1G). With cold exposure, WT (*WT thermoneutral (TN) vs WT cold*) and KO (*KO TN v KO cold*) muscles show a higher expression of metabolic pathways associated with ‘*glucagon*, *AMP-activated protein kinase (AMPK), peroxisome proliferator-activated receptor (PPAR), fatty acid and hypoxia inducible factor 1 (HIF-1) signaling’* as well as ‘*insulin resistance’*. However, the activation of these same pathways is lower in *Actn3* KO muscles, with a reduction in the overall upregulation of the top 7 KEGG pathways observed when accounting for the effect of both genotype and temperature (*Genotype-temperature*, Fig. 1G).

**Figure 1:**
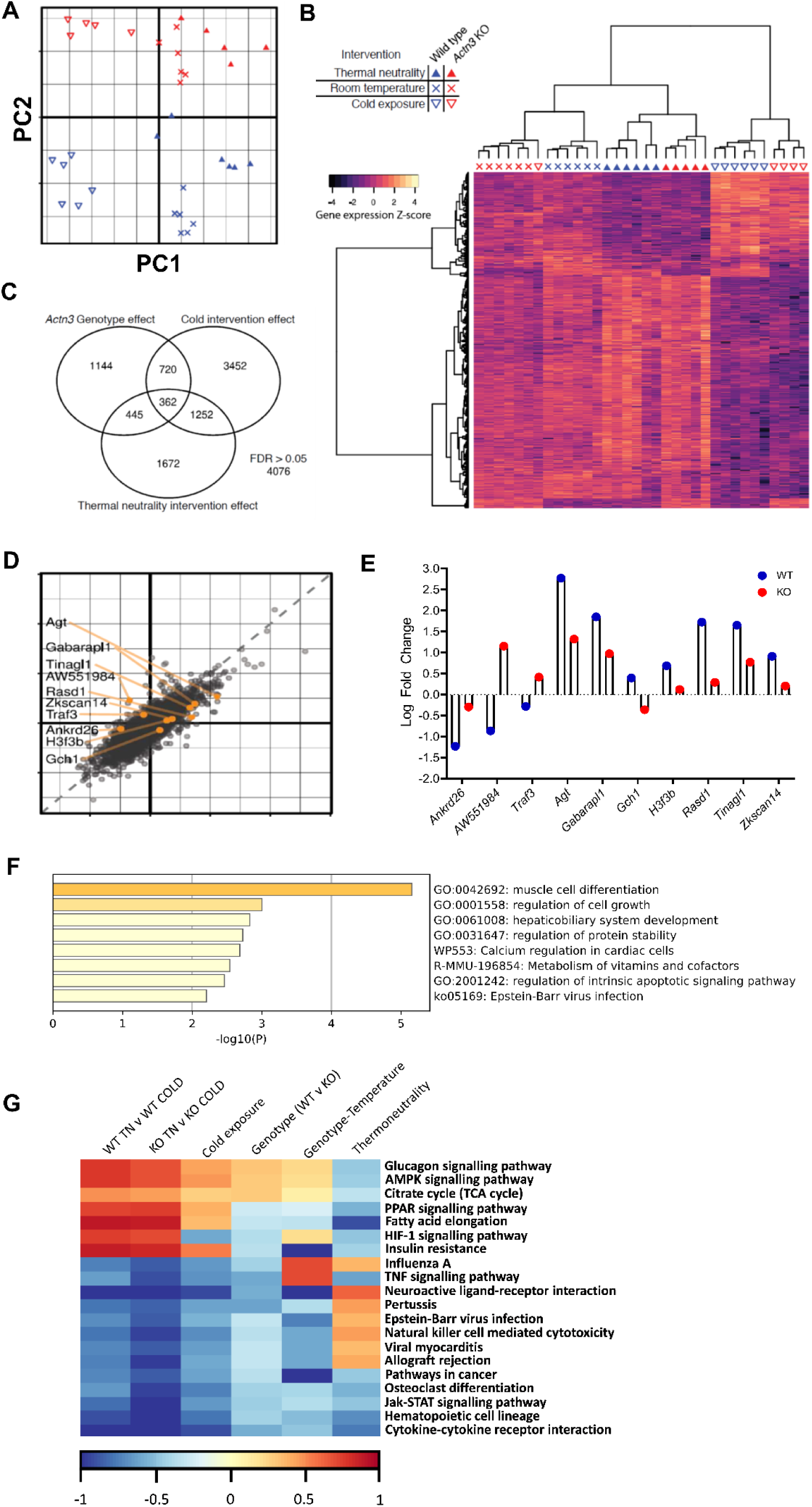
RNA sequencing of skeletal muscle from wild-type (WT) and *Actn3* knockout (KO) mice exposed to thermoneutrality (TN), room temperature (RT) and cold conditions. A) Principal component analysis (PCA) of WT and KO skeletal muscle exposed to either room temperature (RT), cold or thermoneutral (TN) temperature conditions. B) Unbiased hierarchical heat map clustering of all genes. C) Venn diagram of a total of 4076 genes assessed with differences with (genotype), and temperature (cold and thermoneutrality). D) Interaction effect comparing *Actn3* genotype and cold exposure of differentially expressed genes. E) Log fold change of the top ten differentially expressed genes in WT and KO cold exposed skeletal muscle. F) Metascape gene ontology pathway analyses of the significant (adjusted P<0.05) differentially expressed genes in cold exposed KO muscles. G) Kyoto Encyclopedia of Genes and Genomes (KEGG) pathway analyses

### Targeted transcript and protein analyses show that cold-exposed *Actn3* KO muscles have a modified thermogenic response, with lower activation of key Ca^2+^ signaling and mitochondrial activating targets

To further explore the effects of cold exposure on skeletal muscle, we examined an expanded cohort of mice to screen targeted transcripts and protein expression pathways that are modified by *ACTN3* genotype and have been implicated in skeletal muscle thermoregulation.

Following cold exposure, both *Actn3* mRNA and protein abundance are reduced in WT muscles, compared to muscle from WT’s under thermoneutral conditions (mRNA -0.9-fold, protein ∼20%, Fig. 2A). As observed previously, in *Actn3* KO mouse muscles, *Actn3* mRNA and α-actinin-3 protein expression are absent, and there is a compensatory higher expression in the closely related protein, α-actinin-2. *Actn2* mRNA increases (+1.3-fold) in WT and KOs exposed to cold compared to thermoneutral conditions, with *Actn2* mRNA levels higher in KOs than in WT (+1.2-fold) following cold exposure (Fig. 2B). Compared to WT, α-actinin-2 protein abundance is higher in *Actn3* KO muscle at thermoneutrality, (+39%) and following cold exposure (+25%). However, following cold exposure α-actinin-2 was reduced by ∼22% in *Actn3* KO muscle (KO TN *vs*. KO COLD), but this does not occur in WT (WT TN *vs*. WT COLD, Fig. 2B). Together, these data suggest that the skeletal muscle Z-disc, where α-actinin- 2 and -3 co-localize, is altered by acute changes in ambient temperature, and that *ACTN3* genotype influences the Z-disc response to cold exposure.

**Figure 2:**
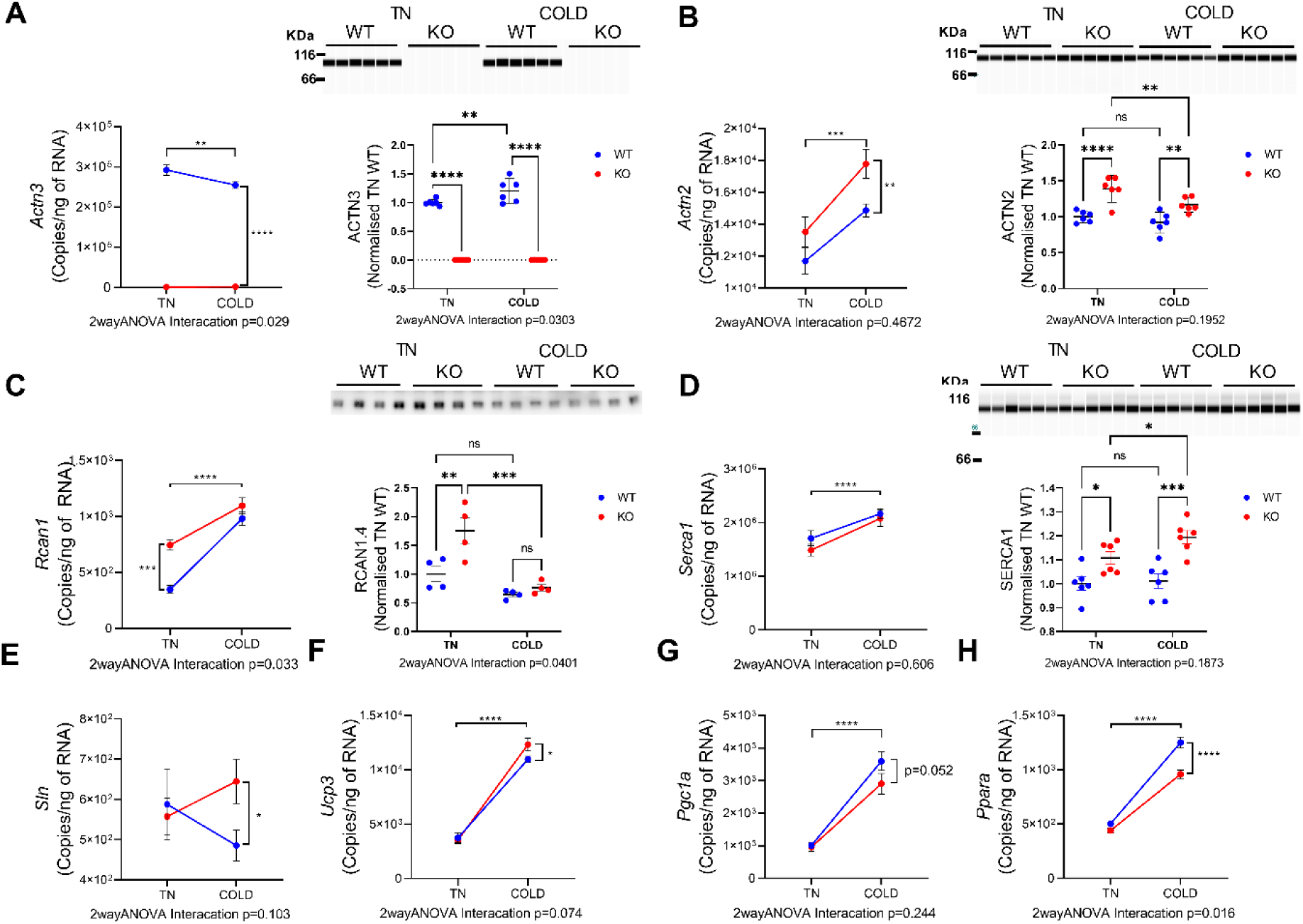
Following acute cold exposure (5 hours at 4°C) targeted transcript and western blot analyses show that *Actn3* KO muscles have an altered Ca^2+^ signaling and thermogenic response compared to wild-type (WT) controls. **A)** *Actn3* mRNA and α-actinin-3 protein expression is absent in *Actn3* KO muscles, and increases following cold exposure in WT muscles. **B)** *Actn2* mRNA and α-actinin-2 protein expression is increased in *Actn3* KO muscles but does not change following exposure to cold **C)** *Rcan1* mRNA and protein expression is increased in KO muscles at thermoneutrality, and is reduced following cold exposure. **D)** *Serca1* mRNA increases in both WT and KO mice following cold exposure. At a protein level, SERCA1 is increased in KO muscles at thermoneutrality and following cold exposure, but remains unchanged in WT mice. **E)** *Sarcolipin (Sln)* mRNA is unchanged in WT and KO mice at thermoneutrality, but increases in KO muscles following cold exposure. **F)** *Ucp3* mRNA is unchanged in WT and KO muscles at thermoneutrality but increases following cold exposure. **G)** *Pgc1α* mRNA is unchanged in WT and KO muscles at thermoneutrality but increases following cold exposure. **H)** *Pparα* mRNA is unchanged in WT and KO muscles at thermoneutrality but increases following cold exposure, with WT mice showing a greater increase than KO’s. N = All values are mean ± SEM. Two-way ANOVA LSD test. *<0.05, **, <0.01, ***<0.001, ****<0.0001

Our previous work has shown that the increase in α-actinin-2 in response to the loss of α-actinin-3, results in an increase in the calcium- and calmodulin-dependent serine/threonine protein phosphatase, calcineurin/Rcan1.4, in both 577 XX human and *Actn3* KO mouse muscles [9]. Here we replicate our initial findings in mice housed at thermoneutrality, with *Actn3* KO muscles showing a 2.1-fold higher *Rcan1* mRNA and Rcan1.4 protein expression compared to WT’s. In addition, cold-exposed WT muscles demonstrate higher levels of *Rcan1* mRNA expression compared to thermoneutral conditions, so that there is no longer a difference between WT and KO muscles following cold exposure (two-way ANOVA p = 0.0326). However, following cold exposure Rcan1.4 protein levels drop in both WT and KO muscles, with a greater decline in KO’s compared to WT’s (two-way ANOVA p = 0.401, Fig. 2D).

Our previous work highlighted a higher expression in the Ca^2+^–sensing protein sarcoplasmic/endoplasmic reticulum calcium ATPase 1 (SERCA1) in *Actn3* KO muscle compared to WT [21]. Increases in SERCA1 and subsequent upregulation of the downstream activator *sarcolipin*, have been previously linked to heat generation in skeletal muscle [25, 26]. Therefore, this upregulation in SERCA1 was hypothesized to be a key mechanism for the proposed positive selection of α-actinin-3–deficient individuals during the migration of modern humans from Africa to the colder Eurasian climates [21]. We therefore examined this pathway in mice housed at both thermoneutrality and following acute cold exposure.

*Serca1* mRNA was increased following cold exposure in both WT and KO muscle (WT +1.3-fold, KO +1.4-fold) (Fig. 2D). A slightly higher SERCA1 protein expression was observed in KO muscles at thermoneutrality (<10%) and to a greater extent following cold exposure (+20%), compared to WT (Fig. 2D). Furthermore, cold-exposed KO’s show a higher sarcolipin (*Sln*) mRNA expression compared to WTs (KO +1.3-fold, Fig. 3E). This provides additional support for alterations in Ca^2+^ handling as a potential contributor to the increase in core body temperature seen in *Actn3* KO and may be implicated in *ACTN3* XX humans following cold exposure [23]. However, SERCA1 or sarcolipin expression differences do not show a significant interaction effect between *Actn3* genotype and temperature by two-way ANOVA, suggesting that the impact of this pathway alone may not explain the improved thermogenic response in *Actn3* KO mice.

**Figure 3:**
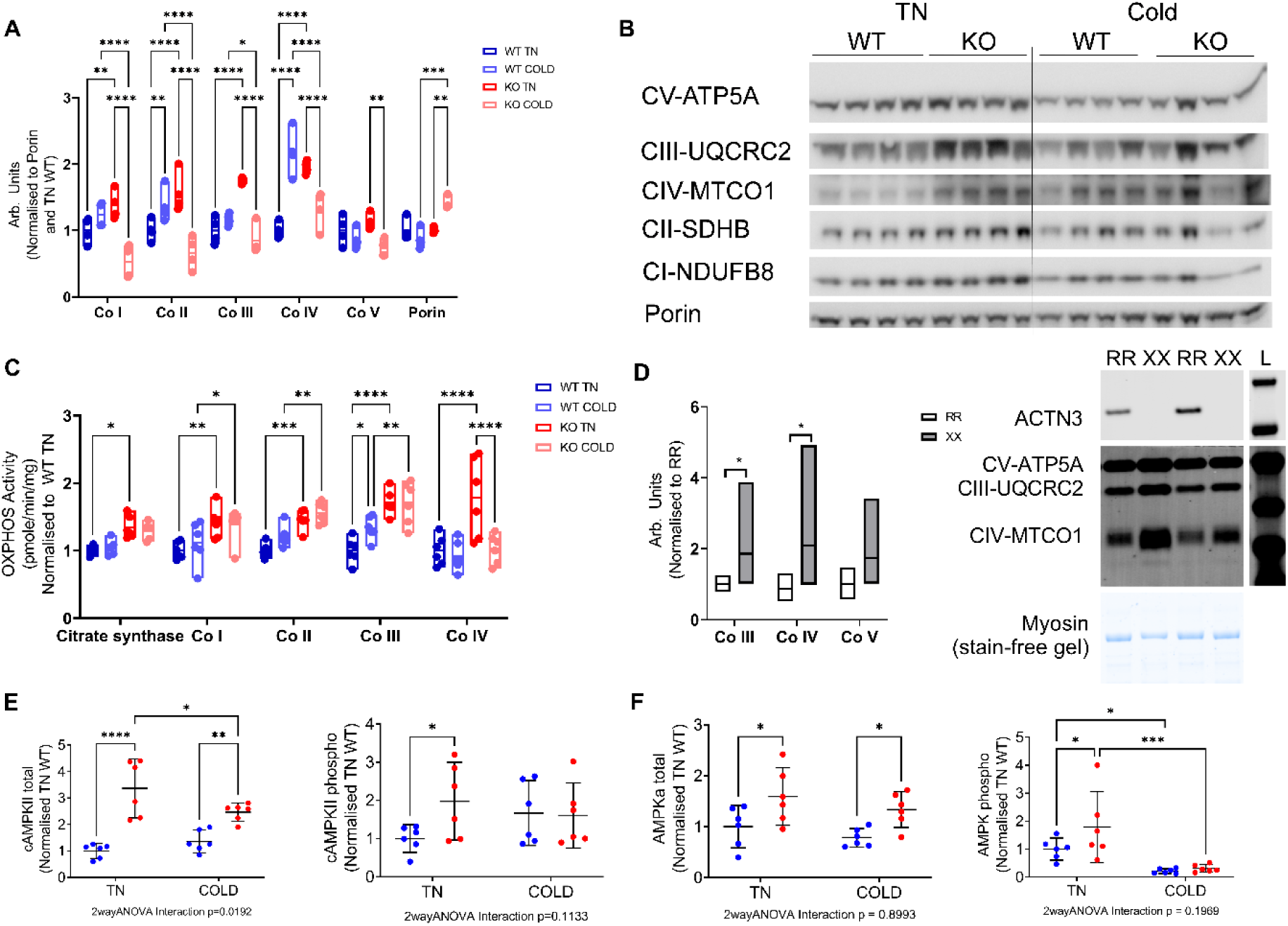
**Mitochondrial OXPHOS and the Ca^2+^/energy sensing proteins Ca^2+^/calmodulin-dependent protein kinase II (CaMKII) and AMP-activated protein kinase (AMPK) are increased at thermoneutrality but reduced following cold exposure in *Actn3* knockout (KO) skeletal muscle.** A) OXPHOS protein expression - complex I (Co I), complex II (Co II), complex III (Co III), complex IV (Co IV), complex V (Co V) and porin abundance in WT and KO muscles at thermoneutrality and following cold exposure. B) Representative western blot image of OXPHOS proteins C) OXPHOS activity assays. Citrate synthase (CS), Co I, Co II, Co III and Co IV in WT and KO muscles at thermoneutrality and following cold exposure. D) Human skeletal muscle OXPHOS Co III, IV and V are increased in XX male individuals collected at thermoneutrality. E) CaMKII total and phosphorylated (phosphor) protein expression in WT and KO muscles. F) AMPK total and phosphorylated (phosphor) protein expression in WT and KO muscles. N = 4 – 6 individuals / genotype. All values are mean ± SEM. Two-way ANOVA LSD test. *<0.05, **<0.01, ***<0.001, ****<0.0001.

Based on these differences we then went on to examine four skeletal muscle thermogenic responsive genes. Increases in uncoupling protein 3 gene (*Ucp3*) expression are considered a primary target for metabolic heat generation in skeletal muscle [27]. In addition, PPARα, PPARγ, and PPAR-coactivator 1α (PGC1α) are well-known transcriptional regulators linked to the expression of metabolic and mitochondrial genes that are upregulated in skeletal muscle following cold exposure [28, 29].

Our data show that *Ucp3* mRNA increases in both WT and KO mice following cold exposure (WT +2.9-fold, KO +3.5-fold). However, cold-exposed KO’s show a trend for a greater increase in *Ucp3* mRNA compared to WT (+1.1-fold, two-way ANOVA interaction p = 0.0743) (Fig. 2F). Similarly, both *Pgc1α* and *Ppar*α mRNA expression are increased following cold exposure in WT and KO mice (*Pgc1α,* WT +3.6 and KO +3.0-fold; *Pparα,* WT +2.5-fold, KO +2.2-fold). Cold-exposed *Actn3* KO mice show lower levels of both *Pgc1α* and *Pparα* mRNA compared to WT mice (*Pgc1α* -0.8-fold, p = 0.052 and *Pparα* -0.8 fold, p <0.001), however this difference was only statistically different by two-way ANOVA for *Pparα* transcripts (Fig. 2G and H).

These data indicate that *Actn3* genotype alters key gene transcripts known to influence skeletal muscle thermogenesis. Taken together these changes suggest an increase in the activation of Ca^2+^–sensitive thermogenic pathways, as well as a lower activation of oxidative metabolism and mitochondrial OXPHOS transcripts in cold-exposed *Actn3* KO muscles compared to WT.

### Mitochondrial/OXPHOS signaling is differentially regulated in *Actn3* KO mice at both thermoneutrality and following acute cold exposure

Based on the specific changes in both gene and protein expression, we went on to explore the effects of cold exposure on mitochondrial OXPHOS protein expression and activity levels to determine if changes in these upstream mitochondrial signaling molecules and pathways differentially modify OXPHOS content and activity in WT and KO mice following an acute exposure to cold conditions (4°C for 5 hours).

Similar to our previous studies [2, 18, 19], compared to WT, KO mice exposed to thermoneutral temperatures (30°C) show higher expression levels of OXPHOS complex I (+∼40%), II (+∼50%), III (+74%) and IV (+95%), but no differences in complex V or in the mitochondrial membrane protein porin (Fig. 3A, B). Similar results were observed in human muscle biopsies collected prior to cold exposure, with muscles from XX individuals showing higher expression levels on both complex III (+50%) and IV (+50%), and no difference in complex V (Fig. 3D).

With cold exposure, WT muscles showed increases in complex II and IV expression, while KO muscles show a reduction in complex I (-72%), II (-71%), III (-32%), and IV (-85%), which is driven by a 54% increase in porin expression (Fig. 3A, B).

Based on the differential OXPHOS protein abundance, we examined the activity of each complex to determine how *Actn3* genotype and the altered OXPHOS protein expression may influence activity. At thermoneutrality, the activity of citrate synthase (CS, +34%), complex I (+42%), II (+46%), III (+70%), and IV (+79%) are all higher in KO compared to WT muscles (Fig. 3J), which is consistent with our protein expression findings.

Following cold exposure, the activity of CS, complex I, II and III remain the same in KOs. However, in WT mice, complex II (+21%) and III (+33%) activity increase. Furthermore, complex IV activity is dramatically reduced in KO muscle following cold exposure (-76%), so that its activity does not show significant differences between WT and KO muscles (Fig. 3C).

This data supports an *Actn3* genotype-specific response in OXPHOS expression and activity following cold exposure, with greater OXPHOS activity in KO mice at thermoneutrality, followed by a significant increase in porin expression with cold exposure.

While this does not result in a significant change in activity of all OXPHOS complexes, complex IV protein expression and activity are significantly reduced in KO muscles following cold exposure.

The striking changes in OXPHOS signaling and activity between WT and *Actn3* KO mice at both thermoneutrality and following cold exposure suggest a key difference in the response to cold. We hypothesized that this differential response could influence important downstream metabolic and signaling pathways associated with energy metabolism. Therefore, we went on to explore the expression of two downstream signaling molecules that are known to be modified by both OXPHOS and Ca^2+^–signaling, calcium calmodulin kinase II (CaMKII) [30] and AMPK.

At thermoneutrality, CaMKII protein expression in KO muscles is remarkably higher than in WT (total +233% and phosphorylated +98%) (Fig. 3E). Following cold exposure, both total and phosphorylated CaMKII decrease in KO muscle, but increase in WT, resulting in no difference in phosphorylated CaMKII expression between genotypes post-cold exposure, with a significant interaction between genotype and temperature observed with total CaMKII expression.

CaMKII is a known activator of the metabolic sensory complex AMPK [31, 32]. A higher abundance of AMPKα (total +59%, and phosphorylated +79%) is observed in KO mice at thermoneutrality (Fig. 4F). However, with cold exposure, total AMPKα remains higher in KO muscles (+55%), while phosphorylated AMPK is dramatically reduced in both KO (- 148%) and WTs (-78.5%) following cold exposure. Together these data reflect the differential activation of key downstream metabolic signaling pathways and the activation of mitochondrial/OXPHOS activity in *Actn3* KO muscles both before and after cold exposure.

**Figure 4:**
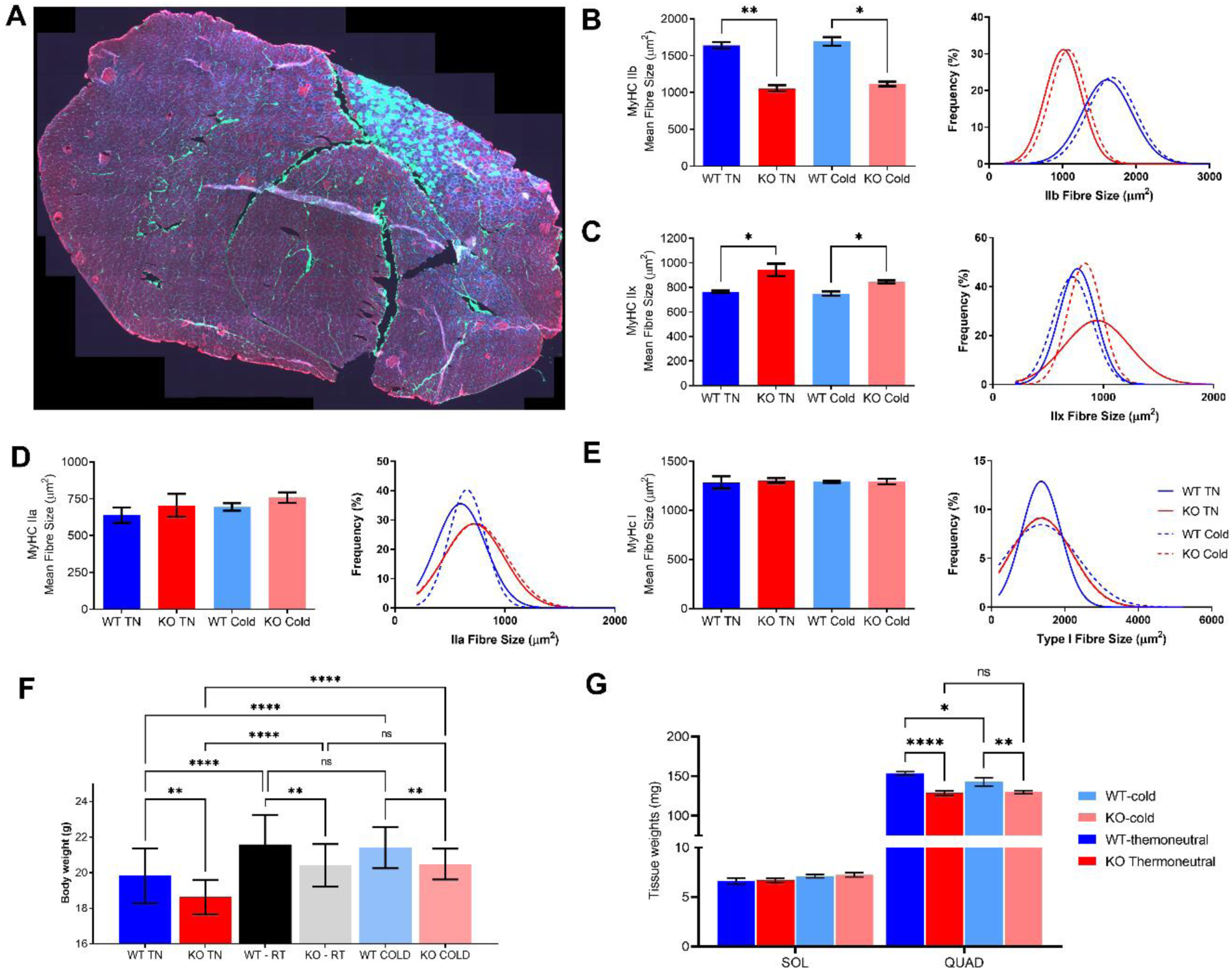
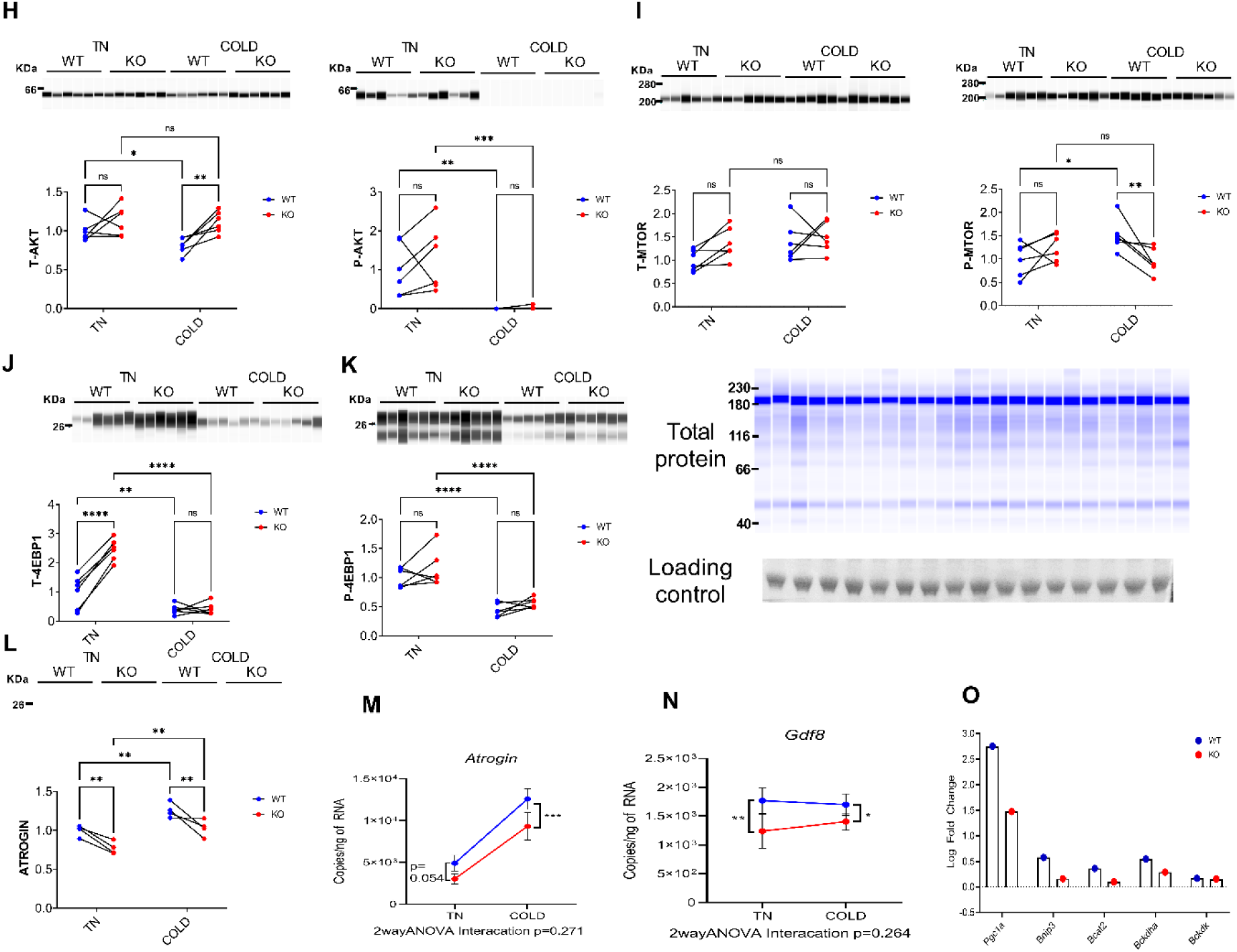
Reduced fast fiber size at thermoneutrality and protection from muscle weight loss following cold exposure is driven by reductions in *atrogin*, *Gdf8* and branch chain amino acid activation in *Actn3* KO muscles A) Representative myosin heavy chain (MyHC) fiber typing image (WT thermoneutral) – Red = IIb, blue = IIa, green = I, black = IIx. B) MyHC type IIb fiber size and frequency C) MyHC type IIx fiber size and frequency D) MyHC type IIa fiber size and frequency E) MyHC type I fiber size and frequency F) Body mass G) Muscle mass – Soleus and quadricep H) AKT – total and phosphorylated I) mTOR – total and phosphorylated J) 4EBP1 – total and phosphorylated K) Total protein for gels A, B, C, D, E, G, Loading control for westerns F L) *Atrogin* mRNA and protein abundance M) *Gdf8* mRNA N) Amino-acid breakdown gene expression Fiber typing and muscle mass analyses N = 6 – 10 mice / genotype / treatment. mRNA transcript analyses N = 12 – 16 mice / genotype / treatment. Protein analyses N = 6 mice / genotype / treatment. All values are mean ± SEM. Two-way ANOVA LSD test. *<0.05, **, <0.01, ***<0.001, ****<0.0001

### Following cold exposure *Actn3* KO muscles show a shift towards a slower muscle phenotype and protection from muscle mass loss due to reduced expression of key protein breakdown/atrophy and amino acid utilization pathway genes

Muscle biopsies collected from *ACTN3* XX humans before cold exposure show an overall lower abundance of fast MyHC muscle isoforms, and a compensatory higher expression of slow type I MyHC protein, compared to *ACTN3* RR. This shift towards a ‘slow’ phenotype would result in an increase in the oxidative capacity in XX muscles and was previously linked to the reduced shivering response and overall improved cold tolerance in XX humans [23]. We therefore performed fiber type analyses of the quadriceps muscle from WT and KO mice to assess MyHC content before and after cold exposure (representative image, Fig. 4A).

Compared to WT mice, *Actn3* KO mice show smaller IIb (Fig. 4B) and larger IIx (Fig. 4C) fiber cross-sectional areas at both thermoneutrality and following acute cold exposure. No differences in MyHC IIa or type I fiber size are observed between genotypes or with acute cold exposure (Fig. 4D, E).

This reduction in fast fiber size in KO muscle has been previously linked to smaller body and skeletal muscle mass in *Actn3* KO mice [17, 18]. *Actn3* KO mice show lower body (Fig. 4F) and skeletal muscle mass (Fig. 4G) at both thermoneutrality and after cold exposure, compared to WT controls. Interestingly, a reduction in quadriceps skeletal muscle mass was observed in WTs following cold exposure whereas no change is seen in *Actn3* KO mice (Fig. 4G, H). This suggests that the loss of α-actinin-3 provides some protection from skeletal muscle breakdown during cold exposure.

Previous work performed in rats has highlighted increases in muscle proteolysis and reductions in protein synthesis during cold exposure [33]. This change in mass was linked to a disruption in the AKT/mammalian target of rapamycin (mTOR) protein signaling pathway, which leads to increases in the muscle specific F-box protein atrogin-1 and increased muscle mass breakdown [33]. Furthermore, our RNA-seq data has highlighted a reduction in the expression of genes associated with protein synthesis and breakdown pathways in cold-exposed *Actn3* KO muscles. Therefore, we went on to explore the impact of cold exposure on skeletal muscle mass and function.

With cold exposure, WT muscles show a significant reduction in total AKT (-30%) compared to no change in KO mice, with the complete absence of phosphorylated AKT in both WT and KO mice after cold exposure (Fig. 5H). Further to this, increased phosphorylated mTOR is observed in WT mice (+51%) after cold exposure, which results in greater mTOR abundance in WT compared to KO skeletal muscle (+55%, Fig. 4I).

**Figure 5:**
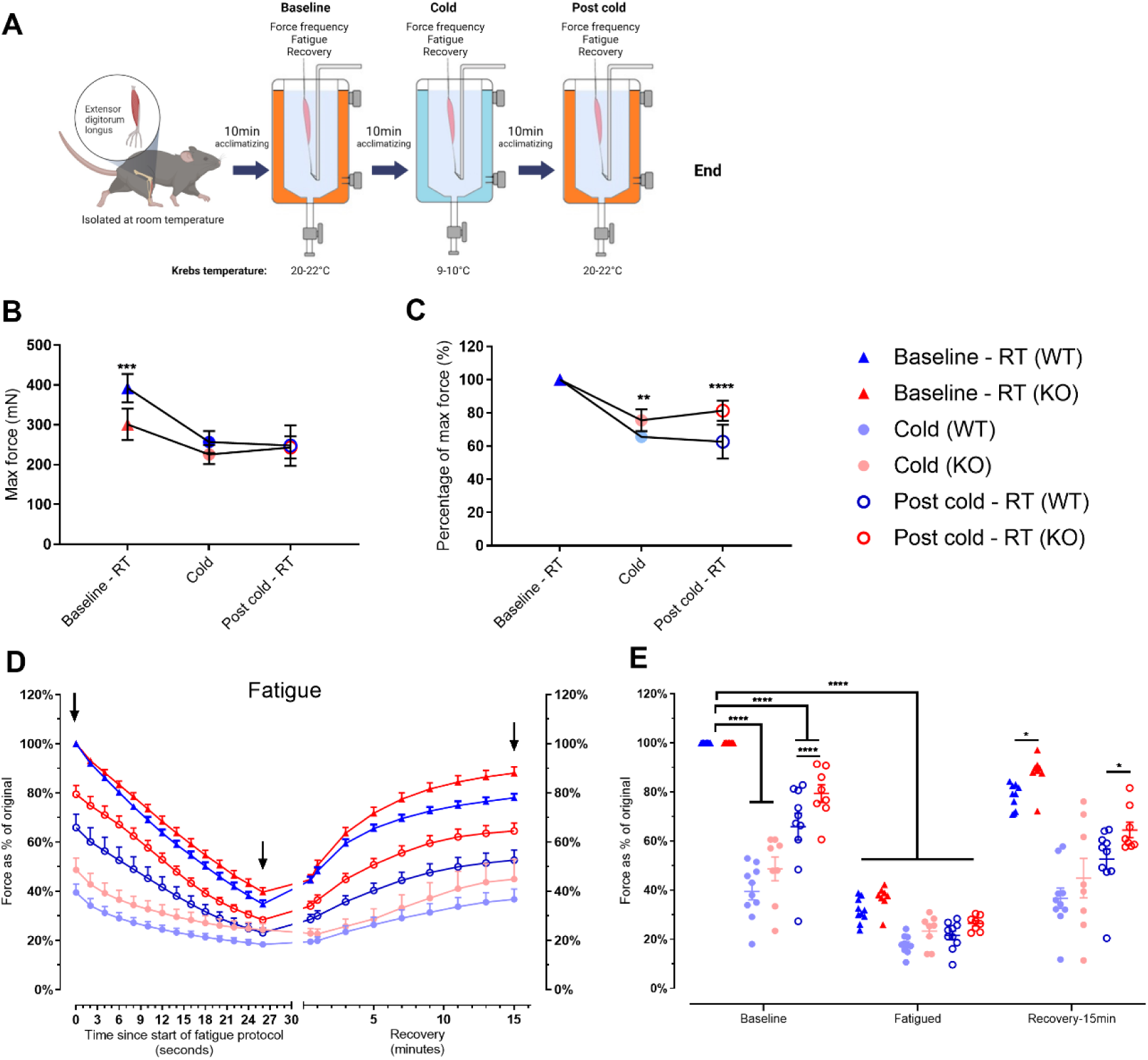
*Ex vivo* extensor digitorum longus (EDL) muscle physiology analyses show reduced force loss and improve fatigue recovery following cold exposure in *Actn3* KO skeletal muscle. A) *Ex vivo* EDL muscle function analyses (force frequency, fatigue and recovery) performed under different krebs solutions temperatures at baseline (20 – 22°C), cold (9-10°C) and post cold (20 – 22°C). The analysis was performed using the same muscle following 10 minutes of acclimatization. B) Absolute maximum force (mN) of WT and KO muscles at Baseline, following cold (9 – 10 °C) exposure and after returning to room temperature (post cold). C) Percentage (%) force loss following cold exposure and return to room temperature (post cold) compared to starting force (Baseline). D) Muscle fatigue and recovery response following incubation at room temperature (Baseline), cold (9-10°C) and post cold (room temperature). Arrows indicate targeted time points shown in panel E. E) Highlight time point analyses at baseline, following fatigue and after 15 minutes of recovery at baseline (solid triangles), following cold exposure (solid circles) and after returning to room temperature (open circles) with individual datapoints for each mouse represented in D. All values are mean ± SEM. One-way ANOVA LSD test. *<0.05, **, <0.01, ***<0.001, ****<0.0001.

Due to the differential response in both AKT and mTOR, we examined the downstream protein synthesis marker initiation factor 4E binding protein 1 (4E-BP1). Increased 4E-BP1 expression is associated with reduced protein synthesis [34]. At thermoneutrality, KO muscles show increased total 4E-BP1 abundance (+189%), which is consistent with our previous findings in mice housed at room temperature [35]. However, following cold, both total and phosphorylated 4E-BP1 are reduced in WT (-61% total; -56% phospho) and KO (-226% total; -59% phospho) muscles, resulting in no major differences between genotypes (Fig. 4J).

Based on these findings we assessed the downstream skeletal muscle atrophy marker E3-ubiquitin ligase, MAFbx/Atrogin-1 and the protein growth/synthesis target growth differentiation factor-8 (*Gdf8*) (also known as myostatin). In KO muscles, both *atrogin-1* mRNA and protein expression are reduced compared to WT at thermoneutrality and following cold exposure (Fig. 4K). Similarly, *Gdf8* mRNA is lower in KO muscles compared to WT at both thermoneutrality and following cold exposure (Fig. 4M). Both atrogin-1 and Gdf8 are known to increase during muscle atrophy [36].

Our RNA-sequencing analysis highlighted altered activation of the branched-chain amino acid (BCAA) breakdown pathway in cold-exposed KO muscles (Fig. 1). This pathway plays a key role in protein synthesis/turnover in skeletal muscle. Reduced activation of genes associated with amino acid turnover in KO muscles could explain the maintenance of muscle mass following cold exposure. We therefore examined the expression of four key BCAA genes, BCL2 interacting protein 3 (*Bnip3*), BCL2 apoptosis regulator (*Bcla2),* and branched chain keto acid dehydrogenase E1 subunit alpha) (*Bckdha)* and branched chain keto acid dehydrogenase kinase (*Bckdk*) in both WT and KO muscles. Here we see a greater activation of *Bnip3, Bcla2* and *Bckdha* gene expression (and no differences for *Bckdk*) in cold exposed WTs compared to KO (Fig. 4N). Previous research has shown that increased expression in these genes results in greater amino acid breakdown and protein turnover in mice [37].

### Loss of α-actinin-3 provides a physiological advantage in muscle performance during and immediately after cold exposure

To further explore the functional implications of these *Actn3* genotype-specific differences, we assessed the *ex vivo* muscle function of isolated *extensor digitorum longus* (EDL) muscles from WT and *Actn3* KO mice exposed to different temperatures (Fig. 5A).

*Ex vivo* muscle function analyses replicated our previous findings [20] and show that KO muscles generate less force under standard temperature conditions (Fig. 5B). With cold exposure, KO muscles lose less force; in fact, max forces during cold exposure and after returning to room temperature did not differ between WT and KO muscles. Consequently, KO muscles produce a greater percentage of muscle strength than WT muscles in the cold and when returned to standard room temperature conditions (Fig. 5C). In addition, KO muscles fatigue less and recover better than WT muscles under normal room temperature conditions (Fig. 5D, E).

These data support a beneficial effect of α-actinin-3 deficiency on the functional attributes of skeletal muscle performance during cold exposure, with *Actn3* KO muscles losing less force, and recovering more quickly, following a fatiguing protocol and acute cold challenge.

### The ACTN3 577X allele (rs1815739) appeared ∼135,000 years ago and has increased in frequency from >42,000 years ago

Using the Genealogical Estimation of Variant Age (GEVA) method [38] we estimate that the age of the X-allele (rs1815739 variant) is 4,650 generations (quality score = 0.92); corresponding to ∼135,000 years before present (BP), given an estimated generation time in humans of 29 years [39] and based on 1 million pairwise haplotype analyses. The distribution of pairwise Time to the Most Recent Common Ancestor (TMRCA) from which the age was estimated (Fig. 6A). We then determined the 95% confidence interval (CI) from the empirical distribution of age estimates, using a haplotype resampling approach, which ranged between 4,480 (130,000 years BP) and 4,830 generations (∼140,000 years BP); the median agreed with our previous estimate of 4,650 generations (Fig. 6B). The rs1815739 X-allele is absent in current archaic human DNA, which suggests that the X-allele originated in *Homo sapiens* and has not been introgressed from Neanderthal or Denisovan hominins [40] (Table S1).

**Figure 6.**
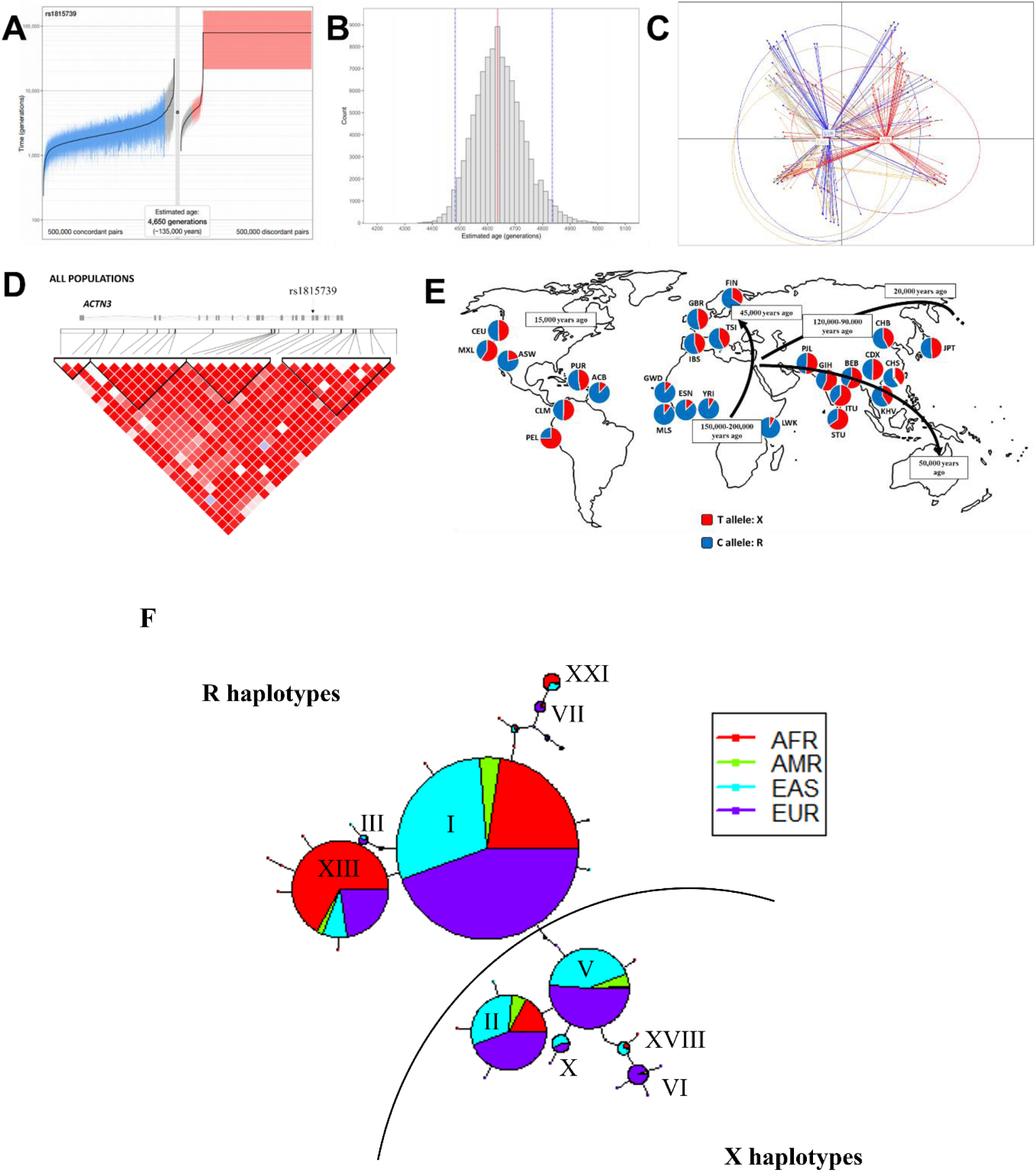
The ACTN3 577X allele originated over 135,000 years ago but its prevalence only increased ∼45,000 years ago with haplotype blocks on chromosome 11 highlighting the unique relationship between X and R alleles in African, American, East Asian and European populations. A) Estimation of variant age of the derived X-allele of the rs1815739 variant in *ACTN3* using the Genealogical Estimation of Variant Age (GEVA) method. B) Histogram showing the distribution of allele age re-estimated through resampling of pairwise time to the most recent common ancestor (TMRCA). C) Principal component analysis for the 577X polymorphisms within European, East Asian, American and African populations shows that the null allele separates based on based on population. D) Linkage disequilibrium (LD) patterns and haplotype blocks for the rs1815739 (577X) variant in *ACTN3*, using published data from the 1000 Genomes project (ftp://ftp-trace.ncbi.nih.gov/1000genomes/ftp/release/20100804/ALL.2of4intersection.20100804.genotypes.vcf.gz). E) Current worldwide distribution of the R (Blue) and X (Red) alleles in the *ACTN3* using 1,000 Genomes Project data [70]. F) Haplotype block on chromosome 11 highlights unique relationships between the *ACTN3* R and X allele in different populations. Abbreviations: AFR, Africa; AMR, Ad Mixed Americans; EAS, East Asia; EUR, Europe.

Following this, we assessed a total of 990 samples – with sample ages of up to 42,000 years BP - from different population cohorts with information on the rs1815739 variant which included, Europe (n = 654 individuals [41–47], Asia (n = 281 [43, 44, 48–51], Central and South America (n = 31 [52]), Oceania (n = 13 [53]) and Africa (n = 11 [54]). Of these, 44.9% of the individuals analyzed carried the X-allele. The only two samples with an age older than 35,000 years BP were X-allele carriers, both located in Europe and Eurasia (Romania, 37,000 to 42,000 years; and Russia, 36,000 to 39.000 years) [42]. We found that 94.1% of the samples (n=17) originating 11,000 to 35,000 years BP carried the ancestral R-allele. While keeping in mind the low sample size of ancient DNA, the frequency of the X-allele in the last 11,000 years was similar to current data, that is, frequencies of 42.5% in samples from Europe and Asia, and of 93.5% in samples from Central and South America. All samples from the Americas corresponded to the period of 0 to 11,000 years BP, with the majority (88%) X-allele positive (**Table S2**).

Results from a discriminant analysis of principal components (DAPC), using populations as clusters, show two selection gradients, between Europe (EUR)/East Asia (EAS), and Africa (AFR). While admixed American (AMR) populations show similarities between both gradients (Fig. 6C). Comparison of data across the four continental populations (EUR, EAS, AMR and AFR) show notable inter-ethnic differences in linkage disequilibrium (LD) patterns and haplotype blocks in EUR and EAS with low LD in AMR or AFR populations (Fig. 6D).

These data support that the X-allele appeared independently from archaic humans (Neanderthal and Denisovan) and provide additional support for higher X-allele frequency recorded with increasing distances from the African continent. All African populations exhibit lower X-allele frequencies (∼10% overall) than the rest of the world, with up to ˃70% in the Americas, and ∼50% in European and Asian populations (Fig. 6E). However, selection for α- actinin-3 deficiency is not as clear in native North American populations, which has been highlighted previously by Amorim et al [15] and Morseburg et al [16,] thereby suggesting that population-specific effects of α-actinin-3 deficiency may occur as a result of genetic drift rather than selection.

To explore the impact of *ACTN3* genotype selection further we examined the relationships among haplotypes using network analysis and identified a haplotype on chromosome 11 from 66,327,673 to 66,329,732 (Fig. 6F). We identified 38 distinct haplotypes in global populations, resulting in 14 577X and 24 577R haplotypes (**Table S3, Table S4, Table S5**). Overall, haplotypes I, II, V, VI, X, XIII and XXI, account for 86.3% of the haplotypes in all populations (Table S4). Of the four haplotypes with frequency ≥0.5% in the region (I, II, V and XIII), the majority of individuals in all populations carry haplotypes with the ancestral C-allele (49.3%, I and XIII haplotypes, vs. 27.4%, II and V haplotypes). The VI and VII haplotypes are completely absent in EAS samples, whereas XIII is present in AFR populations compared to the rest (42.6% vs. 9.1%, 5.1% and 9.4% in AMR, EAS and EUR; respectively, **Table S6**). Phylogenetic reconstruction of the 16 haplotypes with frequency ≥ 0.5% in this genomic region results in a clear separation of haplotypes carrying the ancestral rs1815739-C (577R) and derived rs1815739-T (577X) alleles, which, in combination with previously published work, provides supporting evidence for the selection of α-actinin-3 deficiency in these populations (Fig. 6F). However, this analysis did not assess the impacts of latitude/cold and so further work is required to understand the potential selection mechanisms responsible for the formation of these genetic haplotypes.

## Discussion

In present day humans, the loss of α-actinin-3 is detrimental to sprint and power performance, but the corollary has been reported, with *ACTN3* XX individuals (and *Actn3* KO mice) showing overall improved muscle endurance and greater training adaptability. We have also demonstrated that α-actinin-3–deficient humans and mice show improved heat retention during cold exposure [23].

An increasing body of work has highlighted the importance of skeletal muscle in whole body thermogenesis. Skeletal muscle accounts for ∼40% of an adult human’s body mass and is the major site of glucose uptake and energy expenditure [55]. Skeletal muscle has been confirmed as a key source of whole-body heat generation via both shivering and non-shivering thermogenesis.

In the present study we explore further the known signaling pathways linked to heat production in skeletal muscle to determine the effect of α-actinin-3 deficiency following acute cold exposure.

Firstly, studies in mice have identified muscle-specific mitochondrial uncoupling (proton leak), via sarcolipin-promoted decoupling of the SERCA1 Ca^2+^–sensitive protein pump (Autry et al. 2016), which causes an increase in mitochondrial biogenesis and skeletal muscle oxidative capacity, as a key mechanism for increased heat production following exposure to cold [25, 27, 30, 56–60]. SERCA1 is the primary isoform expressed in fast MHC-containing skeletal muscle fibers in mice. Similar to our previously published data in mice housed at room temperature [21], we now show an increase in SERCA1 protein expression in KO muscles following cold exposure, as well as higher levels of sarcolipin mRNA expression. The increased expression of both SERCA and sarcolipin in cold-exposed KO muscles is relatively small (<20% increase), however considering skeletal muscle accounts for more than 40% of the adult humans’ overall body mass, even a small fraction of heat generated from an increase in the sarcolipin-mediated uncoupling of SERCA could have a significant impact on thermoregulation. Furthermore, the downstream activation of other key proteins and molecules that are impacted by the altered Ca^2+^–SERCA–sarcolipin axis would compound these effects. We therefore went on to explore the *Actn3* genotype-specific changes in downstream Ca^2+^– sensitive targets, including skeletal muscle mitochondrial OXPHOS.

Mitochondrial OXPHOS is one of the most important pathways for cell survival. In eukaryotes, it is responsible for cellular energy production, with alterations in skeletal muscle OXPHOS signaling linked to cold exposure in both mice [61, 62] and humans [63]. Increases in the expression of mitochondrial OXPHOS pathways have been previously observed in *Actn3* KO mice housed at room temperature [2, 19] and a similar network of OXPHOS changes have been linked to the increase in endurance performance and training response in *Actn3* KO mice [2, 9, 17, 18]. Here we show an increase in mitochondrial content (based on an increase in porin expression and CS activity) and reductions in complex IV (content and activity) in KO muscles following cold exposure. These data suggest *Actn3* KO muscles are more efficient at maintaining mitochondrial activity through increased OXPHOS protein expression at thermoneutrality, and that this is preserved following acute cold exposure. In addition the mitochondrial components respond more quickly to acute changes in ambient temperature in KO skeletal muscle compared to WTs. The increase in OXPHOS activity at thermoneutrality in KO mice would result in greater oxidative metabolism, which is similar to the effects seen in chronic cold-exposed mice [56].

Additional downstream markers, Calcineurin (Rcan1.4), CaMKII and AMPK are three key Ca^2+^–sensitive molecules that are all upregulated in KO muscles at thermoneutrality. Following cold exposure, Rcan1.4 and phosphorylated AMPK expression are dramatically reduced in both WT and KO muscles. However, CaMKII (total and phosphorylated) expressions remain higher in KO muscles (at both thermoneutrality and following cold exposure), supporting continued activation of this pathway in KO muscles.

Increases in SERCA-sarcolipin, calcineurin, CaMKII and AMPK are all known to modify mitochondrial/OXPHOS activity. Furthermore, two key mitochondrial transcription factors, *Pgc1α* and *Pparα*, are also differentially expressed in cold-exposed *Actn3* KOs, with reduced activation of both transcripts in KOs after cold exposure.

Taken together these data suggest that α-actinin-3 deficiency results in a “pre-cold acclimatized” muscle phenotype, which results in a faster adaptation to cold exposure and improves the immediate response to acute cold stress in KO muscle.

Skeletal muscle fiber type plays an important role in muscle performance, with fast (type II) muscle fibers generally characterized by powerful contractions and rapid fatigue development, whereas slow-twitch (type I) muscle fibers are fatigue-resistant and used for prolonged activities. α-Actinin-3 is specifically expressed in the fast MyHC-containing skeletal muscle fibers and *ACTN3* genotype is known to modify muscle fiber type proportions, with *ACTN3* 577XX individuals showing a shift towards a slower MyHC profile compared to those that express α-actinin-3 [6, 23]. Here we show a reduction in MyHC IIb fiber size with a compensatory increase in the size of IIx fibers in the predominantly fast quadriceps muscle of

*Actn3* KO mice. Interestingly, similar changes in IIb-IIx fiber sizes have been observed in the skeletal muscle-specific overexpression of PGC1ɑ1 and PGC-1b. Similar to *Actn3* KOs, this shift towards more IIx MyHC was associated with an enhanced oxidative metabolism and improved endurance exercise capacity in PCG1α mice [64, 65].

A benefit of using animal models is that we can assess features that are less accessible in human subjects. One aspect that is difficult to assess in humans is the impact of cold exposure on muscle mass. During cold exposure, energy stores are mobilized because of increased metabolic demand. To supplement this increase in energy needs, skeletal muscle breakdown is increased. At a whole muscle level, WT mice show a greater reduction in muscle mass following cold exposure compared to KO mice, which appear resistant to muscle breakdown and show no change in muscle mass following cold exposure. RNA-seq analyses highlight a reduction in pathways associated with the regulation of *cell growth*, *protein stability* and *metabolism*, as well as *apoptotic signaling* and *energy utilization* in KO muscles, which provides support for the protective effect of α-actinin-3 deficiency on muscle mass pathways following cold exposure.

By assessing the known skeletal muscle protein synthesis/breakdown pathways (AKT, mTOR, Atrogin and Gdf8 and transcripts linked to amino acid utilization (*Bnip3*, *Bcla2* and *Bckdha mRNA*), we show that *Actn3* KO muscles are protected from protein breakdown. Firstly, there were no clear differences in AKT between WT and KO mice at thermoneutrality, however with cold exposure total AKT was reduced in WT muscles but remained unchanged in KO’s. Phosphorylated AKT was unchanged at thermoneutrality but completely absent in both WT and KO muscles following cold exposure. Furthermore, there were no differences in total mTOR between WT and KO mice at thermoneutrality, however following cold exposure, phosphorylated mTOR increased in WT muscles, but remained unchanged in KOs, resulting in an overall increase in mTOR abundance in WT compared to KO muscles. These differences did not correspond to a change in the downstream protein breakdown marker 4EBP1, which suggests that increased protein synthesis is not a key pathway in acute cold exposure KO muscles.

We observed a consistent down-regulation in targets associated with muscle atrophy and amino acid breakdown in KO muscles at both thermoneutrality and following cold exposure. This included lower atrogin-1 protein abundance and transcript, as well as reduced *Gdf8* mRNA expression. Further to this, increased *Pgc1α* expression has been linked to greater branch chain amino-acid breakdown [37]. With cold exposure, we see a clear increase in *Pgc1α* expression in both WT and KO muscles, however *Actn3* KO mice show a reduction in *Pgc1α* mRNA activation following cold. This reduced activation in *Pgc1α* is associated with a similar reduction in the branch chained amino-acid breakdown transcripts *Bnip3*, *Bcla2* and *Bckdha*, in KO muscles, with no differences in the *Pgc1*-α insensitive target *Bckdk*, suggesting that the reduction in *Pgc1α* mRNA activation in KO muscles results in reduced branch chain amino acid signaling in KO muscles.

Since the upregulation of atrogenes is common in various muscle disuse models [36], reduced activation of these targets in KO muscles provides yet another advantage linked to α- actinin-3 deficiency during acute cold exposure. This protection against muscle atrophy stimuli in KO muscles has been reported previously using other models of muscle breakdown, including denervation, immobilization, and glucocorticoid induced muscle atrophy [35, 66]. Similarly, reduced activation of Gdf8 is consistent with increased preservation of muscle mass in α-actinin-3 deficiency [35].

Furthermore, in wild type mice a reduction in circulating Gdf8 has been shown to result in increased muscle OXPHOS, as well as improved skeletal muscle exercise and thermogenic responses via a novel brown adipose tissue (BAT)-skeletal muscle crosstalk mechanism [67]. While we have previously shown that α-actinin-3 deficiency does not modify BAT function during cold exposure [23], the skeletal muscle-specific reduction in Gdf8 seen in *Actn3* KO muscles is associated with features similar to those reported by Kong et al., with improved mitochondrial OXPHOS and greater temperature retention in *Actn3* KO mice. Therefore, the skeletal muscle specific reductions in genes linked to muscle atrophy (Atrogin-1 and *Gdf-8*) and branched chain amino acid (*Pgc1α, Bnip3*, *Bcla2* and *Bckdha mRNA*) pathways in KO’s, together with the preserved expression of protein synthesis/breakdown pathways (AKT and mTOR) are consistent with the reduction in muscle atrophy and altered OXPHOS in KO muscles following cold exposure. Together, these data provide additional support for the beneficial effect of α-actinin-3 deficiency, both before and after cold exposure.

Finally, we explored the functional implications of the broad-reaching molecular effects seen in α-actinin-3 deficiency and cold exposure using *ex vivo* muscle function analyses of isolated EDL muscles exposed to different temperatures. By altering the temperature of the physiological solution from standard room temperature (22°C) down to 9-10°C, we were able to define the impacts of cold exposure on skeletal muscle function and performance. By returning the same solution back to standard room temperature (22°C), we also assessed the muscle’s ability to recover from an acute cold shock.

These analyses demonstrate reductions in muscle strength, increased muscle fatigue and poor fatigue-recovery, in response to cold exposure in WT muscles. However, *Actn3* KO muscles show a greater preservation in muscle strength, improved fatigue response and greater muscle recovery following cold exposure. Similarly, *ex vivo* muscle function analyses suggest that the multi-system changes in KO muscles improve both muscle function and performance following cold exposure.

Further to these mechanistic studies in *Actn3* KO mice we show that the *ACTN3* 577X allele originated in modern humans over 145,000 years ago and has increased in frequency from at least 42,000 years ago. This occurred independent of early human and Neanderthal interactions and is linked in time to modern human migration out of Africa into the European environment.

Both protection from the impacts of cold exposure and improved endurance performance in *ACTN3* 577XX/*Actn3* KO individuals is likely to contribute to improved survival and therefore could account for the increase in *ACTN3* X-allele frequency in modern humans which has been reported previously [13–15]. However, more recent work raises the possibility that the increase in X-allele frequency in non-African populations is not linked to positive selection for cold [16].

An increase in endurance capacity could also provide an evolutionary advantage, and was recently highlighted as a potential selection measure in modern humans [68]. Importantly in skeletal muscle, the mechanisms associated with improved endurance performance and cold tolerance are similar [69], which suggests a potential link between these two phenotypes. However, our work did not explore the potential selection of the *ACTN3* 577X allele with either latitude (cold) or endurance performance and further work is required to correlate this phenotype with any possible selection pressures.

## Conclusion

By comparing mice housed at thermoneutrality (∼30°C), room temperature (∼22°C) and following an acute cold challenge (5 hours at 4°C), we have now defined both the molecular (transcript and protein) and functional (*ex vivo* muscle function) effects of α-actinin-3 deficiency during cold exposure to show that *Actn3* KOs are better able to respond to an acute cold stress. Our comprehensive analysis has shown that the loss of α-actinin-3 modifies key structural (α-actinin-2), Ca^2+^ signalling (SERCA1-sarcolipin, calcineurin (Rcan1.4), CaMKII), mitochondrial (*Pgc1α*, *Pparα*, AMPK and OXPHOS complexes) as well as protein synthesis/breakdown pathways (Atrogin-1, Gdf8 and branched chain amino-acid breakdown) that together provide an overall protective response during cold exposure. This includes an increase in whole body heat retention, reduced muscle mass loss, preserved strength and improved fatigue recovery in *Actn3* KO mice following cold exposure.

Taken together these data provide a plausible molecular and mechanistic explanation for the benefits of the *ACTN3* 577X null allele in modern humans through specific changes in the skeletal muscles response to cold exposure. We also provide data to show physiological changes in muscle function that reinforce the important role that skeletal muscle plays in modifying whole body metabolism and heat production in mammals.

## STAR Methods

### Animal Experiments

This study was approved by The Murdoch Children’s Research Institute (MCRI) Animal Care and Ethics committee (ACEC #760). All experiments were performed using 12-week-old female WT and *Actn3* KO mice on a C57BL/6J genetic background. All mice were maintained in a 12h: 12h cycle of light and dark. Mice were singly housed in cages kept at either 30 °C (thermoneutrality), 22°C (room temperature) or 4 °C (cold) as previously published [71]. Briefly, mice housed at thermoneutrality (30 °C) were acclimatized at this temperature for 20 h (with food and water *ad libitum*) prior to commencement of 5-hour body temperature measurements. Food, water and bedding were removed from cold-exposed mice during the 5- hour cold exposure period. Core body temperature was measured by rectal probe (BAT-12 microprobe thermometer) over a 5-hour period between the times of 0800 and 1400. Temperatures were measured at 0, 30, 60, 90, 120, 180, 240, and 300 min. Body weights were recorded before and after the 5-hour temperature measurement period. At the conclusion of the temperature assessment, all mice were euthanized by cervical dislocation and tissues were collected for further analysis.

### RNA Extractions and Purification

Total RNA was extracted from ∼50mg of mouse quadriceps and ∼20mg of BAT tissues by homogenising and lysing the samples with a T10 basic ultra-turrax disperser (IKA) at 20,500rpm in 1mL of TRIsure solution (Bioline Pty. Ltd, Meridian Biosciences, Australia). After homogenisation, lysates were centrifuged at 12,000rcf for 10 minutes at 4°C to remove cellular debris. The supernatant containing total RNA was transferred into RNAse-free microcentrifuge Eppendorf tubes where 200µl of chloroform was added. The solution was then mixed vigorously by inverting the tubes several times for 15 seconds and left to incubate for 3 minutes at room temperature, followed by centrifugation at 12,000rcf for 15 minutes at 4°C for subsequent phase separation. After centrifugation, the upper layer of aqueous solution containing the RNA phase was transferred without disturbing the interphase to another new RNAse-free microcentrifuge Eppendorf tube with 500µl of isopropanol, and left for 30 minutes at room temperature to precipitate total RNA. For BAT, the fat monolayer was carefully avoided when transferring the upper layer of aqueous solution. Total RNA was then pelleted by centrifugation at 12,000rcf for 10 minutes at 4°C and the supernatant discarded. 1mL of 70% ethanol was next added to wash the RNA pellet by inverting the tubes several times and centrifuging at 7500rcf for 5 minutes at 4°C. The supernatant was discarded and the resultant pellet was air-dried before dissolving and resuspending the pellet in 30µl of Milli-Q water. The resultant RNA solution was then transferred into a new RNAse-free microcentrifuge Eppendorf tube. Using the RNeasy Mini Kit (Qiagen), total RNA was purified as per the company’s protocol and eluted in 30μl of Milli-Q water. RNA integrities (RIN value) and total RNA concentrations were then measured using TapeStation (Agilent Technologies 2200) according to manufacturer’s instructions based on evaluating the ratios of 18S ribosomal RNA and 28S ribosomal RNA subunits. RNA samples were then aliquoted to be stored at -80°C until use for further analyses.

### RNA-sequencing of skeletal muscles from Actn3 KO and WT mice

Skeletal muscle (quadriceps) was collected from *Actn3* WT and KO mice after 5 hours of core body temperature analyses across the three treatment groups (Thermoneutral, TN, Room-temperature, RT, and cold exposed). A total of 18 WT and 16 KO mice underwent RNA sequencing using the Illumina HiSeq 2500 platform as per manufacturers instructions. Raw read data was processed using the Illumina BaseSpace RNA Express application (Illumina Inc. 2016). Briefly, sequencing reads were aligned using STAR ultrafast RNA seq aligner [72] in the SAM file format [73], then counted using HTSeq [74]. The resulting genewise count data was analyzed using the R (3.6.0) statistical programming language (R Core Team 2018). Modelling of differential expression was conducted using the voom precision weights approach [75] in the limma (3.40.6) [76] package from the Bioconductor (3.9) project.

### cDNA Synthesis

Based on initial RNA stock concentrations obtained by TapeStation (Agilent Technologies 2200), samples were diluted with Milli-Q water to a final concentration of 50ng/μl. The diluted RNA samples were quantified using Qubit 3.0 fluorometer (Thermo Fisher Scientific) as per manufacturer’s instructions. All RNA samples were then further diluted to 25ng/μl and 1ng/μl RNA, and 4μl of 25ng/μl RNA or 2μl of 1ng/μl RNA samples (depending on the target gene of interest as per table 2) were reverse transcribed in a final volume of 20μl containing Milli-Q water and reagents from the High-Capacity cDNA Reverse Transcription Kit (Thermo Fisher Scientific) as per the company’s guidelines. Reagents consisted of 10X RT buffer, 10X Random Primers, dNTPs, MultiScribe Reverse Transcriptase and RNAse inhibitor. Samples were next loaded into a thermal cycler (MyCycler, Bio-Rad) to carry out cDNA synthesis with thermal cycling conditions as follows: primer annealing at 25°C for 10 minutes, DNA polymerisation at 37°C for 60 minutes, enzyme deactivation at 95°C for 5 minutes and an infinite hold at 4°C.

### Digital droplet Polymerase Chain Reaction (ddPCR) Droplet Generation and PCR

Droplet digital polymerase chain reaction assay was next conducted, using a PCR reaction assay mixture of 24µl assembled from 2X QX200 ddPCR EvaGreen Supermix, Milli-Q water, 18μM forward primer, 18μM reverse primer (Bio-Rad Laboratories Pty Ltd, Australia), and 0.5-4μl of synthesized cDNA (depending on the target gene of interest as per Table 2) or Milli- Q water as a negative control. The mixture was pipetted into a twin.tec 96-well plate (Bio-Rad) to a final volume of 24µl for lipid droplet generation. The plate was then heat sealed at 180°C for 5 seconds with foil (PX1 Plate Sealer, Bio-Rad), vortexed and centrifuged at 4000rpm for 1 minute before loading into an Auto Droplet Generator (Bio-Rad), where each sample was partitioned into 20,000 uniform sized lipid droplets with the transcripts of interest and cDNA distributed randomly into the droplets. The resultant droplet-partitioned samples were then transferred into a twin.tec 96-well plate (Bio-Rad). The sample plate of droplets was removed from the instrument within 30 minutes of droplet generation completion, heat sealed at 180°C for 5 seconds with pierceable foil (PX1 Plate Sealer, Bio-Rad) and placed in a thermal cycler (T100, Bio-Rad) for subsequent PCR amplification within each droplet. The thermal cycling conditions were as follows: 1 activation cycle of 5 minutes at 95°C, 40 denaturation cycle of 30 seconds at 96°C and annealing cycles of 1 minute at 56-60°C depending on the target gene of interest as per table 2, a post-cycling step of signal stabilization of 5 minutes at 4°C and 5 minutes at 90°C, followed by an infinite 4°C hold, with all cycling steps performed using a 2°C per second ramp rate.

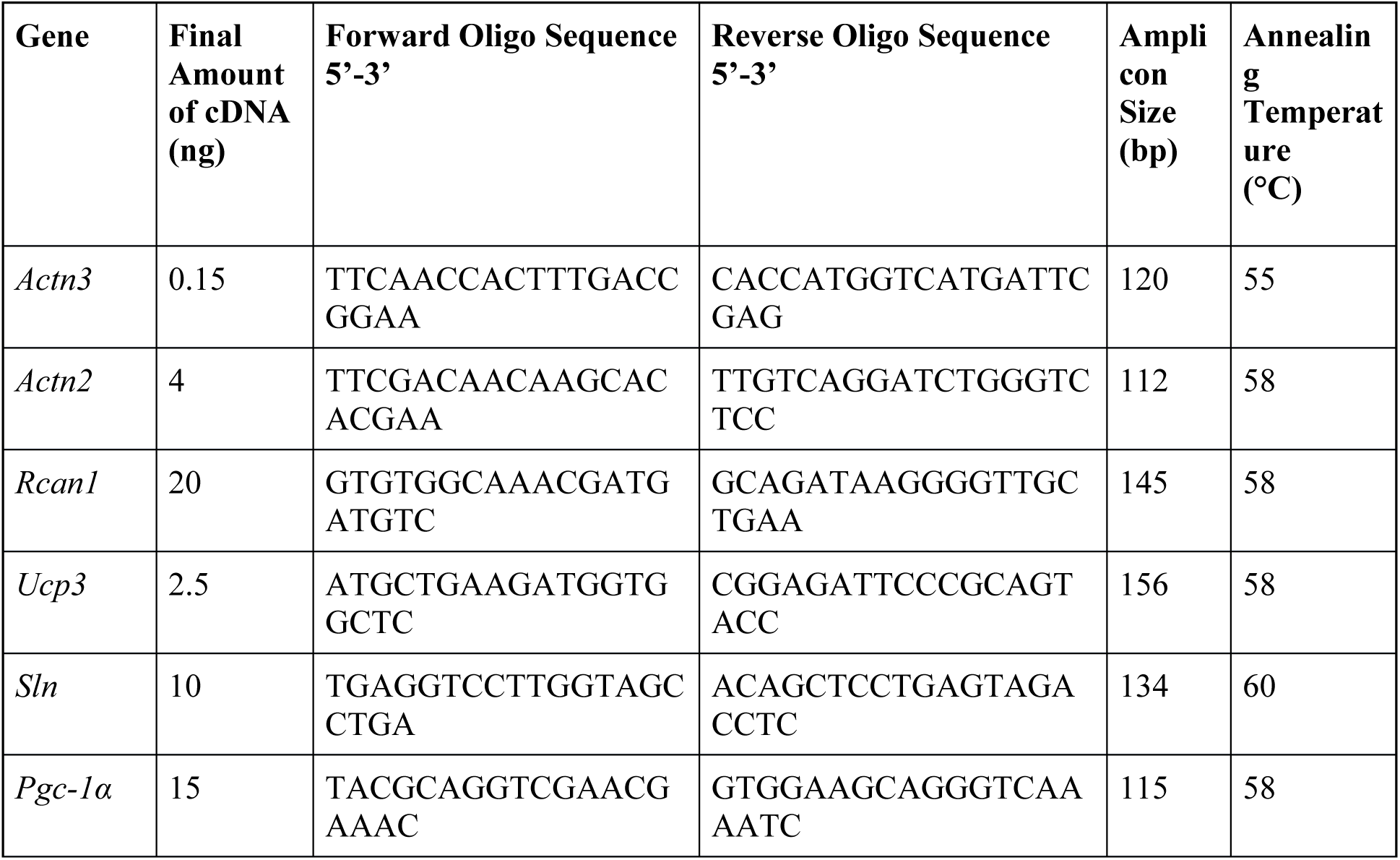
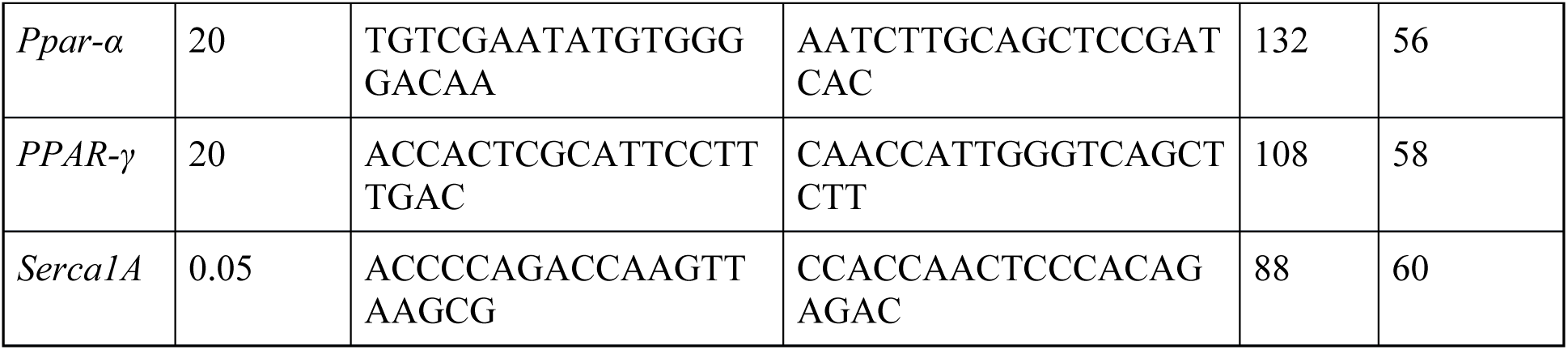

### Droplet Reading and Analyses

Following PCR amplification, the sample plate was loaded on the QX200 Droplet reader (Bio-Rad) and the assay information was entered into the software QuantaSoft (Bio-Rad) that accompanied the droplet reader. Individual droplets from each sample well were read and analysed by the droplet reader in the Evagreen fluorescent channel to give a readout of positive and negative droplets indicating the presence or absence of the target gene respectively. The fraction of positive droplets in a sample were then counted and fitted to a statistical Poisson algorithm to provide plot readouts of absolute quantification and concentration of the transcript of interest by the analysis software QuantaSoft (Bio-Rad). Once results were obtained from the droplet reader, lipid droplets counts were checked using the software QuantaSoft (Bio-Rad) and the rejection criteria for excluding any sample from subsequent analysis included lipid droplet counts lesser than 10,000 for each sample. The 1D amplitude plot of fluorescence amplitude was next visualised to determine data quality based on positive compared to negative droplets separation and droplet scattering. A baseline threshold was applied equally across all samples depending on the fluorescence amplitude to distinguish positive from negative droplets, and readings of the target were obtained in copies of RNA transcript/μl of reaction and copies of RNA transcript per reaction. Data was exported as a comma separated value (.csv) files and further analyzed using Microsoft Excel, where readings of the target obtained in RNA transcript copies/μl of reaction and RNA transcript copies per reaction were adjusted for RNA loading and converted to absolute values of RNA transcript copies/ng of RNA. All data was entered into GraphPad Prism 9 for statistical analyses. Table 2 shows a list of target genes of interest used for DDPCR.

### Western Blot

Quadriceps muscles from WT and KO mice were frozen in liquid nitrogen, powdered and then lysed by sonication in 4% SDS lysis buffer containing 65mM Tris pH 6.8, 4% SDS, 1:500 Protease Inhibitor Cocktail solution (Sigma-Aldrich, Australia), 1X PhosSTOP Phosphatase Inhibitor Cocktail solution (Sigma-Aldrich) and Milli-Q water. Total protein concentrations were then assessed using Direct Detect (Thermo Fisher Scientific, Australia) as per manufacturer’s instructions. For western blot, 20µL of protein lysates containing a total of 10µg (for RCAN1) or 20µg (for OXPHOS and Porin) of protein, 50mM DTT, 10% glycerol and bromophenol blue dye were then mixed and heated at 50°C for 5 minutes before loading onto Invitrogen Bolt 4-12% Bis-Tris Plus precast gels (Thermo Fisher Scientific) in 1X Bolt MOPS SDS running buffer (Thermo Fisher Scientific) in a mini gel tank (Thermo Fisher Scientific) for SDS-PAGE separation at 200V for 30 minutes. Proteins were then transferred to polyvinylidene fluoride membranes (PVDF, Merck Millipore) in transfer buffer containing 1X Tris-Glycine (Bio-Rad), 0.075% SDS, 20% methanol and MilliQ-water in a Trans-Blot Cell (Bio-Rad) at 100V for 30 minutes at 4°C. The membranes were then blocked with 5% skim milk in 1× TBST for 1 hour at room temperature, washed with 1X TBST for 5 minutes 3 times, and probed with primary antibodies diluted in 5% skim milk TBST using 1:1000 OXPHOS Rodent WB Antibody Cocktail (Sapphire Bioscience, Australia), 1:2000 Anti-

VDAC1/Porin [20B12AF2] (ab14734, Abcam, United Kingdom) and 1:500 RCAN1 (Anti- DSCR1 C-terminal, D6694, Sigma-Aldrich) overnight at 4°C. Blots were then washed in 1X TBST for 5 minutes 3 times, probed with 1:10,000 respective secondary antibodies diluted in 5% skim milk TBST for 1 hour at room temperature and washed in 1X TBST for 5 minutes 3 times. Membranes were then developed with ECL reagents (Amersham Biosciences, United Kingdom) and imaged on Image Quant (GE Healthcare, Australia). Densitometry was performed using ImageJ image processing software (NIH) and quantified using the area under the curve.

For semi-automated western blotting, 0.1mg/mL (for total protein), 0.02mg/mL (for α-actinin- 3 and α-actinin-2) or 0.2mg/mL (for SERCA1) of protein lysates, primary antibodies including 1:3000 α-actinin-3 (ab68204, Abcam), 1:2500 α-actinin-2 (ab68167, Abcam) and 1:5000 SERCA1 (ab109899, Abcam), and their respective secondary antibodies were prepared and added into a 12-230 KDa 25 capillary cartridge separation module (Protein Simple) as per manufacturer’s instructions and loaded onto the WES machine (Protein Simple) for western blotting. Densitometry measures were performed using Compass software (Protein Simple). All results were then normalized to total protein and presented relative to WT thermoneutral controls.

### Mitochondrial activity assays

Spectrophotometric enzyme assays assessing mitochondrial oxidative phosphorylation (OXPHOS) and citrate synthase activities were performed in duplicate using homogenized WT and *ACTN3* KO tibialis anterior muscles. OXPHOS enzyme assays were performed on postnuclear supernatants as described previously by Frazier et al [77]. Briefly, post-nuclear supernatants were prepared at ∼4°C by homogenizing 50 mg (wet weight) of tissue in 9 volumes of Tissue buffer (5 mM HEPES, 1 mM EGTA, 210 mM mannitol, 70 mM sucrose, pH 7.2) with 10 strokes in a glass–glass homogenizer, followed by centrifugation at 600 RCF for 10 min. Enzyme assays measured the change of absorbance in a 1 mL cuvette at 30°C over 3 min in 50 mM potassium phosphate buffer, pH 7.4 using a Cary 300 Bio Spectrophotometer (Agilent Technologies Australia). Protein levels were measured using bicinchoninic acid (BCA) and enzyme activities were expressed as initial rates (nmol/min/mg) except for complexes III and IV, which were expressed as rate constants (/min/mg). Enzyme activities were expressed as a mean, normalized to WT thermoneutral control.

### Ex vivo muscle physiology

All experiments were conducted blinded to genotype.

The EDL muscles of WT (n = 10) and *Actn3* KO (n = 8) mice housed under standard room temperature conditions (22°C) were dissected from the hindlimb and tied by their tendons to a dual force transducer/linear tissue puller (300 Muscle Lever; Aurora Scientific Instruments, Canada) using 6-0 silk sutures (Pearsalls, United Kingdom). During dissection the muscle was kept in a temperature controlled organ bath containing room temperature (22°C) Krebs solution with composition (in mM): 4.75 KCl, 118 NaCl, 1.18 KH2PO4, 1.18 MgSO4, 24.8 NaHCO3, 2.5 CaCl2 and 10 glucose, 0.1% fetal calf serum and bubbled continuously with carbogen to maintain pH at 7.4.

The EDL muscle was stimulated to contract by delivering a current between two parallel platinum electrodes, using an electrical stimulator (701C stimulator; Aurora Scientific Instruments). All contractile procedures were designed, measured, and analyzed using the 615A Dynamic Muscle Control and Analysis software (DMC version 5.417 and DMA version 5.201; Aurora Scientific Instruments). At the start of the experiment, the muscle was set to optimal length (Lo), which produced maximal twitch force. The initial supramaximal stimulus was given at 1 ms, 125 Hz for 1 s, and force produced was recorded as Po, the maximum force output of the muscle at Lo.

Each muscle then underwent three sets of consecutive contractile procedures, with the temperature of the Krebs solution varying after each protocol as outlined in Figure 5A. The protocol started with baseline measurements at room temperature (20 – 22°C), followed by a 10 minute acclimatisation in cold (9-10°C) Krebs solution when the second muscle physiology protocol was performed, and concluded with another 10 minute acclimatisation in Krebs solution returned to room temperature (20 – 22°C) to complete the post-cold exposure measurements. Each contractile procedure included a maximum force reading, a fatigue protocol, and a 15-minute recovery period using methods as previously published [78].

Briefly, force-frequency curves were generated before fatigue and after recovery to measure muscle contractile function as previously published . Trains of stimuli given at 1 ms at different frequencies, including 2, 15, 25, 37.5, 50, 75, 100 and 125 Hz for 1 s for EDL and the force produced was measured. A 30-s rest occurred between each frequency.

To test the rate of fatigue development, trains of stimuli were given at 1 ms, 125 Hz for 1 s for a total of 15 were given with 1 second break after each stimuli, upon completion muscle recovery was measured by delivering the same stimuli at 30 s, 1 min, 3 min, 5 min, 7 min, 9 min, 11 min, 13 min and 15 min respectively. Force values were expressed as a percentage of first fatigue contraction. Whole muscle length was measured using vernier muscle calipers while sitting at optimal length within the organ bath. All muscles were blotted dry (Whatmans filter paper DE81 grade) and weighed using an analytical balance (GR Series analytical electronic balance) following contractile procedures. Forces were normalized with respect to an estimate of physiological cross-sectional area, according to the equation CSA=MM/(Lo*D), where MM is the muscle mass, Lo is the optimal length, and D is the density of skeletal muscle (1.06 g/cm3), to enable comparisons between muscles of differing sizes and weights.

### The ACTN3 rs1815739 - X allele frequency in modern humans

The worldwide distribution of the ACTN3 rs1815739 SNP was estimated using the frequencies recorded in the 1,000 Genomes Project, which contains data of 2,504 individuals from 26 populations [70].

We accessed the 1000 Genomes genetic data of 629 individuals from different ethnic backgrounds (ftp://ftp-trace.ncbi.nih.gov/1000genomes/ftp/release/20100804/ALL.2of4intersection.20100804.genot <u>ypes.vcf.gz</u>). The core haplotype surrounding rs1815739 was identified using biallelic markers across a region surrounding ACTN3 (Chr11:66,313,866-66,330,805, hg19, 37 SNPs) in haploview [79, 80], from all 26 populations in the 1000 Genomes data. Pairwise fixation index (FST) for discriminant analysis of principal components (DAPC) analysis for all pairs of populations were obtained using the adegenet and hierFstat R packages [81, 82]. To reduce complexity or noise, haplotypes from each population were combined revealing a haplotype block defined on Chromosome 11 from position (hg19) 66,327,673 to 66,329,732 with a high linkage disequilibrium (LD) pattern (Figure S1). Haplotypes within the defined haplotype block were constructed into networks with the Pegas R package [83]. We inferred the phylogeny of the haplotypes using the ape and ggtree packages [83, 84].

#### The rs1815739-X allele and modern genomes

We investigated putative associations between rs1815739 SNP and other phenotypes using the atlas of genome wide scan association studies (GWAS), which contains data from 4,756 GWAS across 3,302 unique traits (https://atlas.ctglab.nl) [85] and the Electronic Health Record-Derived PheWAS codes from the Michigan Genomics Initiative [86]. We also assessed effects on gene expression using Regulome DB annotations [87] (https://regulomedb.org/regulome-search) and GTEx data [88] (https://gtexportal.org/home/).

We analyzed the worldwide distribution of the rs1815739 single nucleotide polymorphism (SNP) using the modern human data of the frequencies as recorded in the 1,000 Genomes Project [70], which contains data of 2,504 individuals from 26 populations: *African ancestry:* African Caribbean people in Barbados (ACB), Americans of African Ancestry in South West (United States of America, ASW), Esan in Nigeria (ESN), Gambian in Western Divisions (GWD), Luhya in Webuye (Kenya, LWK), Mende in Sierra Leone (MSL), and Yoruba in Ibadan (Nigeria, YRI); *American ancestry:* Colombians from Medellin (CLM), Mexican Ancestry from Los Angeles (MXL), Peruvians from Lima (PEL), and Puerto Ricans from Puerto Rico (PUR); *East Asian ancestry*: Chinese Dai in Xishuangbanna (CDX), Han Chinese in Beijing (CHB), Southern Han Chinese (CHS), Kinh in Ho Chi Minh City, Vietnam (KHV) and Japanese in Tokyo (JPT); *European ancestry:* Utah Residents with Northern and Western European Ancestry (CEU), Finnish in Finland (FIN), British in England and Scotland (GBR), Iberian Population in Spain (IBS) and TSI Tuscany in Italy (TSI); and *South Asian ancestry:* Bengali from Bangladesh (BEB), Gujarati Indian from Houston, Texas (GIH), Indian Telugu from the United Kigndom (ITU), Punjabi from Lahore, Pakistan (PJL), and Sri Lankan Tamil from the United Kindom (STU).

We accessed the 1000 Genomes genetic data of 629 individuals from different ethnic backgrounds (ftp://ftp-trace.ncbi.nih.gov/1000genomes/ftp/release/20100804/ALL.2of4intersection.20100804.genot <u>ypes.vcf.gz</u>). We then performed haplotype analysis for AFR, AMR, EAS and EUR superpopulations across a +-5 kb extended region surrounding *ACTN3*, using Haploview version 4.1 software (https://www. broadinstitute.org/haploview/haploview) (43 single nucleotided polymorphisms [SNPs])[79].

#### Early and extinct Homo genomes

In order to investigate the human evolutionary history of the rs1815739, we checked the genotypes for rs1815739 in available data from extinct Homo lineages, that is, Neanderthals (n=4) and Denisovans (n=2) [89–91], as well as from an ancient human, Ust’-Ishim (*i.e.*, the ∼45,000-year-old remains one of the early modern humans to inhabit western Siberia)[92]. We determined archaic-specific missense variants in *ACTN3* from Neanderthal and Denisovan individuals, evaluating their presence in present-day humans. The coding sequence of α- actinin-3 protein was extracted from the publicly available exome database (http://cdna.eva.mpg.de/neandertal/exomes/VCF/)[90]. The sequences were aligned to the reference Homo sapiens protein sequences in Uniprot (http://www.uniprot.org/).

We also analyzed ancient DNA data extracted from the datasets of David Reich’s group (https://reich.hms.harvard.edu/), coming from different studies published in the last years, with samples from various locations: Europe [41–47], Asia [43, 44, 48–51], Central and South America [52], Oceania [53] and Africa [54]. The designation of T (X) or C (R) allele for rs1815739 in each sample was extracted using PLINK software (v1.9)[93].

#### Origin of the X-allele of rs1815739

We estimated the time of origin of the derived X-allele of the rs1815739 variation, using the Genealogical Estimation of Variant Age (GEVA) method [38]. The age of an allele is estimated based on the distribution of TMRCA inferred between pairs of haplotypes, where time (generations) is modelled jointly from mutational and recombinational information from sequencing data (GEVA joint clock model). Our analysis used data from the 1000 Genomes Project [94], where the X-allele is found at a sample-wide frequency of 40%, and we used Ne=10,000 and a mutation rate of µ=1.2x10^-8^ as scaling parameters, as well as data from HapMap2 to infer the recombination rate along the genome [95]. We estimated the age of the X-allele using one million haplotype pairs (*i.e.* 0.5 million concordant pairs, since both haplotypes carry the focal allele) and 0.5 million discordant pairs (one carrier and one non-carrier haplotype), after correctly mapping alleles by their ancestral and derived states (see below).

Note that GEVA assumes that the reference allele represents the ancestral allelic state, but which, according to information available through Ensembl, was not the case for the rs1815739 variant in the 1000 Genomes data set (see https://human.genome.dating/snp/rs1815739). We therefore extended the method to optionally consider external information about the ancestral allelic state of the variants present in a given data set; the software is available online: https://github.com/pkalbers/geva/tree/ancallele. Here we used information from Ensembl (release 95) to map the ancestral and derived allelic states for all variants on Chromosome 11 in 1000 Genomes data, because GEVA considers all variants observed along the sequence in the inference of TMRCA at a given locus.

### Statistics

Statistical analyses are described in the figure legend of each analysis. Briefly, all analyses were performed using GraphPad Prism v9.1. Differences equal to or lower than the p-value of 0.05 were considered significant. Data shown are either scatter plots or summary data with mean values and error bars indicating either SEM or 95% confidence interval (CI) as outlined in the figure legends.

