## Supplemental Tables for "α-Actinin-3 deficiency protects from the effects of acute cold exposure through altered skeletal muscle Ca^2+^ and OXPHOS signaling"

### Supplementary TABLES

**Table S1**. Analysis of the rs1815739 variant in the α-actinin-3 (*ACTN3*) gene in data from extinct *Homo* species.

| Sample | A | C | G | T |
|---|---|---|---|---|
| Sidron[1] | 0.0 | 7.3 | 0.0 | 0.0 |
| Vindija[2] | 0.0 | 30.7 | 0.0 | 0.0 |
| AltaiNea[3] | 0.0 | 26.2 | 0.0 | 0.0 |
| Denisova[4] | 0.0 | 15.6 | 0.0 | 2.0 |
| Mez1[5] | 0.0 | 2.0 | 0.0 | 0.0 |
| DenisovaPinky[6] | 0.0 | 17.7 | 0.0 | 1.0 |
| UST[7] | 0.0 | 13.2 | 0.0 | 0.0 |

Abbreviation: SNP, single nucleotide polymorphism.

[1] http://cdna.eva.mpg.de/neandertal/exomes/VCF/Sidron/
[2] http://cdna.eva.mpg.de/neandertal/exomes/VCF/Vindija/
[3] http://cdna.eva.mpg.de/neandertal/altai/AltaiNeandertal/VCF/
[4] http://cdna.eva.mpg.de/neandertal/Vindija/VCF/Denisova/
[5] http://cdna.eva.mpg.de/neandertal/Vindija/VCF/Mez1/
[6] http://cdna.eva.mpg.de/neandertal/altai/Denisovan/
[7] http://cdna.eva.mpg.de/ust-ishim/VCF/

**Table S2**. Frequency of X (T) and R (C) alleles of the rs1815739 variant in the α-actinin-3 gene (*ACTN3*) gene, as identified in ancient DNA data sets of cohorts in Europe (N=654), Asia (N=281), Central and South America (N=31), Oceania (N=13) and Africa (N=11).

| | Europe | | Asia | | Central and South America | | Oceania | | Africa | |
|---|---|---|---|---|---|---|---|---|---|---|
| **Time period (years BP)** | **X-allele n (%)** | **R-allele n (%)** | **X-allele n (%)** | **R-allele n (%)** | **X-allele n (%)** | **R-allele n (%)** | **X-allele n (%)** | **R-allele n (%)** | **X-allele n (%)** | **R-allele n (%)** |
| 42.000–37.000 | 1 (100%) | 0 (0%) | | | | | | | | |
| 39.000–36.000 | 1 (100%) | 0 (0%) | | | | | | | | |
| 35.000–34.000 | 0 (0%) | 2 (100%) | | | | | | | | |
| 31.000–29.000 | 0 (0%) | 1 (100%) | | | | | | | | |
| 30.000–29.000 | 0 (0%) | 1(100%) | | | | | | | | |
| 28.000–27.000 | 1 (50%) | 1 (50%) | | | | | | | | |
| 19.000–18.000 | 0 (0%) | 2 (100%) | | | | | | | | |
| 17.000–16.000 | 0 (0%) | 1(100%) | | | | | | | | |
| 14.000–13.000 | 0 (0%) | 7 (100%) | 0 (0%) | 1 (100%) | | | | | | |
| 12.000–11.000 | 0 (0%) | 1 (100%) | | | | | | | | |
| 11.000–10.000 | 5 (50%) | 5 (50%) | 2 (67%) | 1 (33%) | 1 (100%) | 0 (0%) | | | | |
| 10.000–9.000 | 3 (20%) | 12 (80%) | 3 (50%) | 3 (50%) | 6 (86%) | 1 (14%) | | | | |
| 9.000–8.000 | 23 (47%) | 26 (53%) | 0 (0%) | 2 (100%) | 2 (100%) | 0 (0%) | | | | |
| 8.000–7.000 | 39 (33%) | 80 (67%) | 2 (50%) | 2 (50%) | 3 (75%) | 1 (25%) | | | | |
| 7.000–6.000 | 20 (40%) | 30 (60%) | 1 (33%) | 2 (67%) | 3 (100%) | 0 (0%) | | | | |
| 6.000–5.000 | 37 (45%) | 45 (55%) | 12 (34%) | 23 (66%) | 1 (100%) | 0 (0%) | | | 0 (0%) | 1 (100%) |

| | | | | | | | | | | |
|---|---|---|---|---|---|---|---|---|---|---|
| 5.000–4.000 | 103 (47%) | 114 (53%) | 18 (44%) | 23 (56%) | 2 (100%) | 0 (0%) | | | 1 (100%) | 0 (0%) |
| 4.000–3.000 | 43 (51%) | 42 (49%) | 63 (46%) | 74 (54%) | 2 (100%) | 0 (0%) | | | | |
| 3.000–2.000 | 2 (25%) | 6 (75%) | 30 (64%) | 17 (36%) | | | 1 (25%) | 3 (75%) | 0 (0%) | 3 (100%) |
| 2.000–1.000 | | | | | | | 0 (0%) | 2 (100%) | 0 (0%) | 3 (100%) |
| 1.000–1 | | | 1 (33%) | 2 (67%) | 9 (100%) | 0 (0%) | 4 (57%) | 3 (43%) | 0 (0%) | 2 (100%) |

**Table S3**. Haplotypes described

| **Haplotypes** | **rs1671064** | **rs2229455** | **rs1815739** | **rs618838** | **rs2229456** | **rs2290463** | **rs607736** | **rs77239910** |
|---|---|---|---|---|---|---|---|---|
| I | A | A | C | C | A | G | G | G |
| II | G | A | T | T | A | G | A | G |
| III | A | A | C | C | A | C | A | G |
| IV | A | G | C | C | C | C | G | A |
| IX | A | G | C | C | C | G | G | A |
| V | G | A | T | T | A | G | G | G |
| VI | G | G | T | T | C | G | G | A |
| VII | A | G | C | C | C | G | A | A |
| VIII | A | G | C | C | C | G | G | G |
| X | G | A | T | T | A | C | G | G |
| XI | A | A | C | C | A | C | G | G |
| XII | A | G | C | C | C | C | G | G |
| XIII | A | A | C | C | A | G | A | G |

| | | | | | | | | |
|---|---|---|---|---|---|---|---|---|
| XIV | G | A | C | C | A | G | G | G |
| XIX | G | A | T | T | A | G | A | A |
| XV | G | A | C | T | A | C | G | G |
| XVI | G | A | T | T | C | G | G | G |
| XVII | G | G | T | T | C | G | A | A |
| XVIII | G | G | T | T | C | G | G | G |
| XX | A | A | T | C | A | G | G | G |
| XXI | A | G | C | C | C | G | A | G |
| XXII | G | G | C | T | C | G | G | A |
| XXIII | A | A | C | T | A | G | G | G |
| XXIV | G | G | T | T | A | G | G | A |
| XXIX | G | G | T | T | A | G | G | G |
| XXV | A | A | C | C | C | G | A | G |
| XXVI | A | A | T | C | A | C | A | G |
| XXVII | G | G | C | C | A | G | A | G |
| XXVIII | G | G | C | C | C | G | G | G |
| XXX | A | A | C | C | C | G | G | G |
| XXXI | G | G | T | C | C | G | G | G |
| XXXII | A | G | C | C | A | G | G | G |
| XXXIII | A | G | C | C | A | G | A | G |
| XXXIV | A | A | C | T | A | G | A | G |
| XXXV | A | A | C | C | A | G | A | A |
| XXXVI | A | A | T | T | A | G | A | G |
| XXXVII | G | A | C | T | A | G | G | G |
| XXXVIII | G | A | C | T | A | G | A | G |

**Table S4**. Major R haplotype frequencies as a percentage of total R haplotypes.

| | **Haplotypes** | | | | | |
|---|---|---|---|---|---|---|
| **Superpopulation** | **I** | **XIII** | **XXI** | **VII** | **III** | **Others** |
| AFR | 11.68 | 18.27 | 3.17 | 0.38 | 0.13 | 3.93 |
| AMR | 1.90 | 0.51 | 0.00 | 0.13 | 0.00 | 0.00 |
| EAS | 14.97 | 2.28 | 1.52 | 0.00 | 0.76 | 2.92 |
| EUR | 22.84 | 6.22 | 0.13 | 2.54 | 1.65 | 4.06 |

Abbreviations: AFR, Africa; AMR, Ad Mixed Americans; EAS, East Asia; EUR, Europe.

**Table S5**. Major X haplotype frequencies as a percentage of total X haplotypes.

| | **Haplotypes** | | | | | |
|---|---|---|---|---|---|---|
| **Super population** | **II** | **V** | **VI** | **X** | **XVIII** | **Others** |
| AFR | 6.17 | 0.21 | 0.00 | 0.00 | 1.28 | 1.28 |
| AMR | 2.34 | 2.13 | 0.64 | 0.00 | 0.00 | 0.00 |
| EAS | 11.06 | 16.17 | 0.00 | 4.89 | 4.47 | 1.06 |
| EUR | 15.96 | 19.36 | 8.72 | 3.40 | 0.21 | 0.64 |

Abbreviations: AFR, Africa; AMR, Ad Mixed Americans; EAS, East Asia; EUR, Europe.

**Table S6**. Haplotype frequencies in the different super populations

| Haplotypes | AFR (2N=338) | AMR (2N=44) | EAS (2N=354) | EUR (2N=522) | TOTAL (2N=1,258) |
|---|---|---|---|---|---|
| I | 27.22 | 34.09 | 33.33 | 34.48 | 32.19 |
| II | 8.58 | 25.00 | 14.69 | 14.37 | 13.28 |
| III | 0.30 | 0.00 | 1.69 | 2.49 | 1.59 |
| IV | 0.00 | 0.00 | 0.28 | 1.72 | 0.79 |
| IX | 0.00 | 0.00 | 0.00 | 1.72 | 0.72 |
| V | 0.30 | 22.73 | 21.47 | 17.43 | 14.15 |
| VI | 0.00 | 6.82 | 0.00 | 7.85 | 3.50 |
| VII | 0.89 | 2.27 | 0.00 | 3.83 | 1.91 |
| VIII | 1.48 | 0.00 | 2.82 | 0.19 | 1.27 |
| X | 0.00 | 0.00 | 6.50 | 3.07 | 3.10 |
| XI | 0.00 | 0.00 | 1.13 | 0.77 | 0.64 |
| XII | 0.00 | 0.00 | 1.69 | 0.38 | 0.64 |
| XIII | 42.60 | 9.09 | 5.08 | 9.39 | 17.09 |
| XIV | 0.89 | 0.00 | 0.28 | 0.19 | 0.40 |
| XIX | 0.00 | 0.00 | 0.28 | 0.00 | 0.08 |
| XV | 0.00 | 0.00 | 0.00 | 0.19 | 0.08 |
| XVI | 0.30 | 0.00 | 0.00 | 0.19 | 0.16 |
| XVII | 0.00 | 0.00 | 0.00 | 0.19 | 0.08 |
| XVIII | 1.78 | 0.00 | 5.93 | 0.19 | 2.23 |
| XX | 0.00 | 0.00 | 0.85 | 0.00 | 0.24 |
| XXI | 7.40 | 0.00 | 3.39 | 0.19 | 3.02 |
| XXII | 0.00 | 0.00 | 0.00 | 0.19 | 0.08 |
| XXIII | 0.00 | 0.00 | 0.28 | 0.38 | 0.24 |

| | | | | | |
|---|---|---|---|---|---|
| XXIV | 0.00 | 0.00 | 0.00 | 0.19 | 0.08 |
| XXIX | 0.89 | 0.00 | 0.00 | 0.00 | 0.24 |
| XXV | 2.37 | 0.00 | 0.00 | 0.00 | 0.64 |
| XXVI | 0.00 | 0.00 | 0.28 | 0.00 | 0.08 |
| XXVII | 1.48 | 0.00 | 0.00 | 0.00 | 0.40 |
| XXVIII | 0.30 | 0.00 | 0.00 | 0.00 | 0.08 |
| XXX | 1.18 | 0.00 | 0.00 | 0.00 | 0.32 |
| XXXI | 0.30 | 0.00 | 0.00 | 0.00 | 0.08 |
| XXXII | 0.30 | 0.00 | 0.00 | 0.00 | 0.08 |
| XXXIII | 0.59 | 0.00 | 0.00 | 0.00 | 0.16 |
| XXXIV | 0.30 | 0.00 | 0.00 | 0.00 | 0.08 |
| XXXV | 0.30 | 0.00 | 0.00 | 0.00 | 0.08 |
| XXXVI | 0.30 | 0.00 | 0.00 | 0.00 | 0.08 |
| XXXVII | 0.00 | 0.00 | 0.00 | 0.19 | 0.08 |
| XXXVIII | 0.00 | 0.00 | 0.00 | 0.19 | 0.08 |
